# A hardware/software system for electrophysiology “supersessions” in marmosets

**DOI:** 10.1101/2020.08.09.243279

**Authors:** Jens-Oliver Muthmann, Aaron J. Levi, Hayden C. Carney, Alexander C. Huk

## Abstract

We introduce a straightforward, robust method for recording and analyzing spiking activity over timeframes longer than a single session, with primary application to the marmoset (*Callithrix jacchus*). Although in theory the marmoset’s smooth brain allows for broad deployment of powerful tools in primate cortex, in practice marmosets do not typically engage in long experimental sessions akin to rhesus monkeys. This potentially limits their value for detailed, quantitative neurophysiological study. Here we describe chronically-implanted arrays with a 3D arrangement of electrodes yielding stable single and multi-unit responses, and an analytic method for creating “supersessions” combining that array data across multiple experiments. We could match units across different recording sessions over several weeks, demonstrating the feasibility of pooling data over sessions. This could be a key tool for extending the viability of marmosets for dissecting neural computations in primate cortex.

## Introduction

The marmoset has drawn attention as a complementary nonhuman primate model system for visual neuroscience. While the dominant primate model system in neuroscience, the rhesus monkey (*Macaca mulatta*), has the advantage of (relatively) rich cognitive abilities, a large body and robust physiology, and an aggressive work ethic, their large and convoluted (gyrified) brains currently limit the number of techniques that can be applied for measurements of neural activity. Thus, despite their excellent trainability for complex tasks and willingness to engage in lengthy experimental sessions, the scale and variety of neurophysiological questions that can be addressed have been somewhat limited by practical constraints. Recently, the common marmoset (*Callithrix jacchus*) has emerged as a complementary primate model system because of their smooth (lissencephalic) cortex, opening up a much larger number of cortical areas to the use of large-scale chronically implanted electrode arrays (in addition to other techniques). However, a major current concern for adopting the awake behaving marmoset for detailed quantitative studies is their tendency to perform far fewer trials per session compared to macaques. Such a behavioral limitation would result in correspondingly smaller amounts of neural data (and hence, statistical power) per experiment, undercutting the other advantages of the species, and likely limiting their applicability as a powerful neurophysiological complement to the sorts of quantitative neuroscience work done in macaques.

To redress this fundamental potential limitation, we have developed a straightforward, user-friendly tool for recording from large-scale arrays in marmosets while surmounting the relatively short behavioral sessions performed by this smaller (and more delicate) species. First, we report successful long-term electrophysiological recordings using a new type of multi-electrode array for which primate use has not yet been reported in publication to our knowledge, but which is commercially available. These “3D” arrays are available with customizable electrode spacing not just across a 2D grid, but also along the depth of individual shanks. The arrays yielded good quality single-unit (SUA) and multi-unit (MUA) activity, as demonstrated in two different marmoset cortical areas (area MT, and the posterior parietal cortex, PPC). Second, we introduce a transparent means for identifying activity recorded on these arrays, not just within individual sessions, but — importantly — *across* sessions. This integration of hardware and software solutions allowed for data from the same unit to be combined over multiple behavioral sessions, into what we termed “supersessions.” This brings the statistical power of awake-behaving marmoset neurophysiology closer to that of macaques on a per-unit basis, while still allowing for larger scale recordings and/or powerful complementary tools, such as patch-clamp and optogenetics, that are more challenging to perform in macaques.

Here, we describe both the physiological and computational components of this tool and demonstrate its potential usefulness for transcending the behavioral limitations of marmosets into the realm of detailed, quantitative assessments of neural activity at large scales. Furthermore, the tool we introduce here is intentionally straightforward, meaning it can be readily implemented by others, as well as extended when ongoing updates to hardware and software emerge. We conclude by describing current limitations and how updates to this tool could further improve it.

To provide a bit more detail before delving into the results, we found that implanting commercially-available 3D “N-form arrays” (ModularBionics, Berkeley, CA, USA) resulted in high quality, stable unit activity in marmosets. In our hands and experiences, this reflected a significant step forward in neural recording success, as two prior attempts using more common types of 2D planar arrays (Utah, Black rock systems) yielded lower-quality outcomes (one successful insertion without detectable spikes and one with spiking activity for about three months after implantation). Although our goal was simply to record neural activity and not to mechanistically understand why a particular array style works better or worse, our hypothesis is that there is a reduced initial damage due to the lower number of shanks of the N-form array, allowing to avoid vasculature and permitting a slow insertion style. In contrast to single shanks and arrays with a single row of shanks, we believe that long-term stability is improved by a better fixation of the brain tissue, reducing chronic respiratory micromotion (***Prodanov and Delbeke, 2016***), while eventually compromising a smaller brain volume for blood circulation than the larger 2D planar arrays.

Given the success of the neural recording hardware in yielding qualitatively impressive neural activity over long time periods, we asked whether such recordings would yield a broad sample of neurons that change from experiment to experiment or if they would yield longer recordings of the same neurons. In the first case, we could ask how neural responses generalize across the population, but would overestimate generalization if we recorded from the a substantial subset of neurons from day to day, but did not recognize that in our analyses. In the second case, we could obtain longer recordings for individual units and hence a higher statistical power. We thus designed a method to systematically compare and match (distributions of) spike waveforms across sessions. Our method identifies units from individual sessions independently, and then integrates spike clusters from new recordings into known, existing ones identified in prior sessions. Analyses of units can therefore be performed over multiple experimental sessions.

In order to achieve a representation of spike shapes that was robust to potentially varying noise levels and/or forms across experimental sessions, we extracted simple properties of spike shapes in a narrow window around their peak. This was achieved by matching a family of predefined templates on a GPU to yield a parametric representation of local excursions in the raw voltage traces, which included conventional unit spiking activity, spike events from weaker or more distant neural sources, and noise. Unit isolation was performed as a multivariate classification problem, similar to conventional approaches (***Pachitariu et al., 2016***; ***Rossant et al., 2016***; ***Chung et al., 2017***; ***Hilgen et al., 2017***; ***Jun et al., 2017a***; ***Lee et al., 2017***; ***Chaure et al., 2018***; ***Diggelmann et al., 2018***; ***Yger et al., 2018***). In our method, we did not threshold spikes during a detection step, but clustered shapes of local minima in the voltage traces. The resulting clusters were then matched across recording sessions. Although we are not deeply attached to this particular spike sorting approach, we provide it as a robust, intuitive starting point, which we validated against a more sophisticated and complex spike-sorting package. Its simplicity also allows for online views of sorting results during experiments, which could be useful for experimental decisions even if more sophisticated sorting routines are employed post hoc.

Finally, in addition to laying out the hardware and software that allows for supersession-style electrophysiology in marmosets with chronic recording arrays, we also provide starting-point quantifications of the performance of this system. These metrics confirm the applicability of this system to many conventional neurophysiological experiments given the performance level that arises from the current arrays and implantation style, as well as the spike sorting algorithm. However, the greater value of these metrics is in future use, as they will allow for comparisons of relative performance (in matters such as falsely-matched units across sessions) as array technology changes, as surgical procedures are refined, and as different spike sorting algorithms are applied.

Taken together, this work puts forth a synthesis of commercially-available hardware and intuitive software that allows experimenters to overcome one of the major limitations of the marmoset as a model species by introducing the concept of supersessions. More generally, this framework may support better integration of work done in marmosets and macaques, allowing these two awake-behaving primate preparations to have greater scientific overlap and thus to more solidly allow for their relative strengths and weaknesses to be considered.

## Results

### Neural activity apparent for more than 9 months on chronically-implanted 3D arrays

We recorded single and multi-unit (hereafter, “unit”) activity in the brains of 2 marmosets, one with a 3D N-form array in and around the middle temporal area (MT), the other with an identical array placed in posterior parietal cortex (PPC). For both arrays (Figure 1 A, B, respectively), we were able to record spiking activity starting a week after insertion. Activity lasted for a duration of at least 9 months, as depicted in Figure 1 (top rows). Figure 1 (second rows) show, in comparison, the relatively short durations of individual recording sessions (approximately a half hour to an hour). These durations likely reflect a lower bound on how long marmosets will work, as they were largely determined by the animal’s preponent motivation to engage in various visual tasks with no fluid or food restriction.

**Figure 1.**
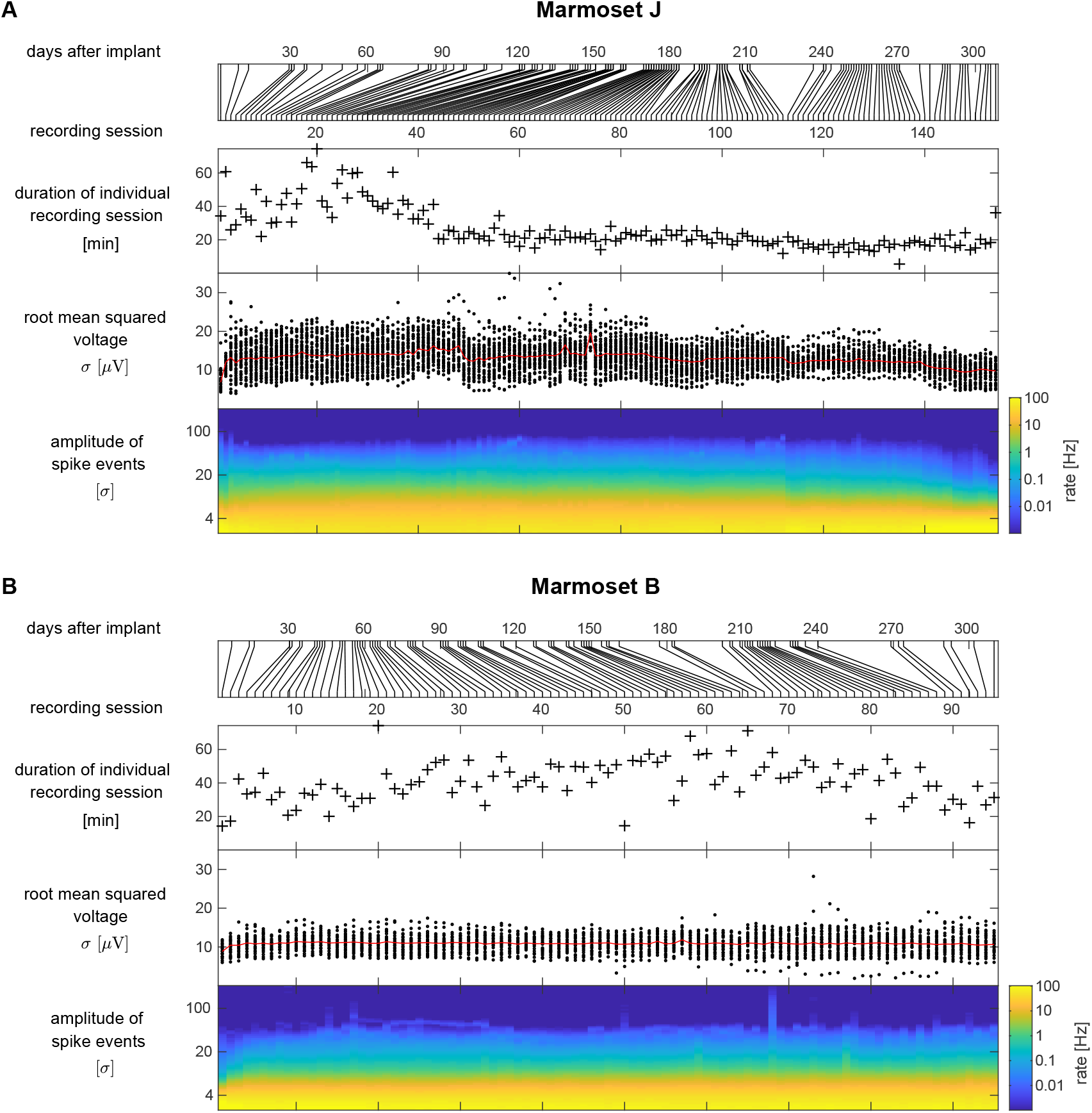
Long-term stability of arrays. (**A**) marmoset J. Top panel: Illustration when individual recording sessions were performed. For clarity, the plots below and in subsequent Figures reflect individual recording sessions rather than time. Second row: Durations of electrophysiological recordings in individual sessions. Third row: Root-mean-squared voltage fluctuations of the common averaged, 300 Hz high-pass filtered data (scatter plots for active electrodes, average shown in red). Bottom row: Amplitude histograms of detected events, averaged across electrodes. (**B**) Same statistics for marmoset B. **Figure 1–source data 1**. Source data to generate this Figure

Signal amplitudes (Figure 1, third rows) were fairly constant over long periods of time, perhaps with the first two weeks after implantation yielding smaller signals before stabilizing (i.e., first few recording sessions, visible at the very left of the plots). A gradual decline in signal amplitude was further apparent after about 7 months for marmoset J. Detected events (see Methods) had a wide amplitude range of relatively sparse (0.1 – 10 Hz) events, indicative of spiking activity (Figure 1, bottom rows). Taken together, these descriptions of the behavior of the animals and the signals from the electrode arrays lay the groundwork for attempting to stitch together data from multiple, subsequent recording sessions. The next critical step would be identifying unit activity that could conservatively be identified across such sessions.

### Spike clusters overlap in consecutive sessions

Our goal was to identify spikes from the same units across recording sessions. This required measures that would be robust to noise, in the sense that spikes from other neurons would not perturb or distort characterization and identification of a given unit. To that aim, we focused our analysis on a very short temporal window, including only the depolarization phase of a spike, represented by a local minimum in the raw voltage traces.

For each local minimum (i.e., putative spike) in the raw voltage trace, we determined: (a) amplitude, measured as the dot product with a template (of unit power), expressed in standard deviations (*σ*), as calculated on the high-pass filtered voltage traces; (b) width, measured as the full width at half minimum; and (c) symmetry, measured as the ratio of its falling and rising phase durations (i.e., a 1 : 2 ratio means that recovering back to baseline took twice as long as reaching the voltage minimum).

These parameterized shape characterizations of the units were put into 3D-histograms (marginals shown in Figure 2 A) for each recording session, and clustered using a watershed algorithm (see Methods for details). This procedure yielded shape clusters (cyan markers in Figure 2 A) for every session in a common coordinate system to allow for cross-session comparisons of spike shapes. Shape clusters between consecutive sessions often looked very similar, and so we further tested whether they likely reflected spikes from the same or from different units.

**Figure 2.**
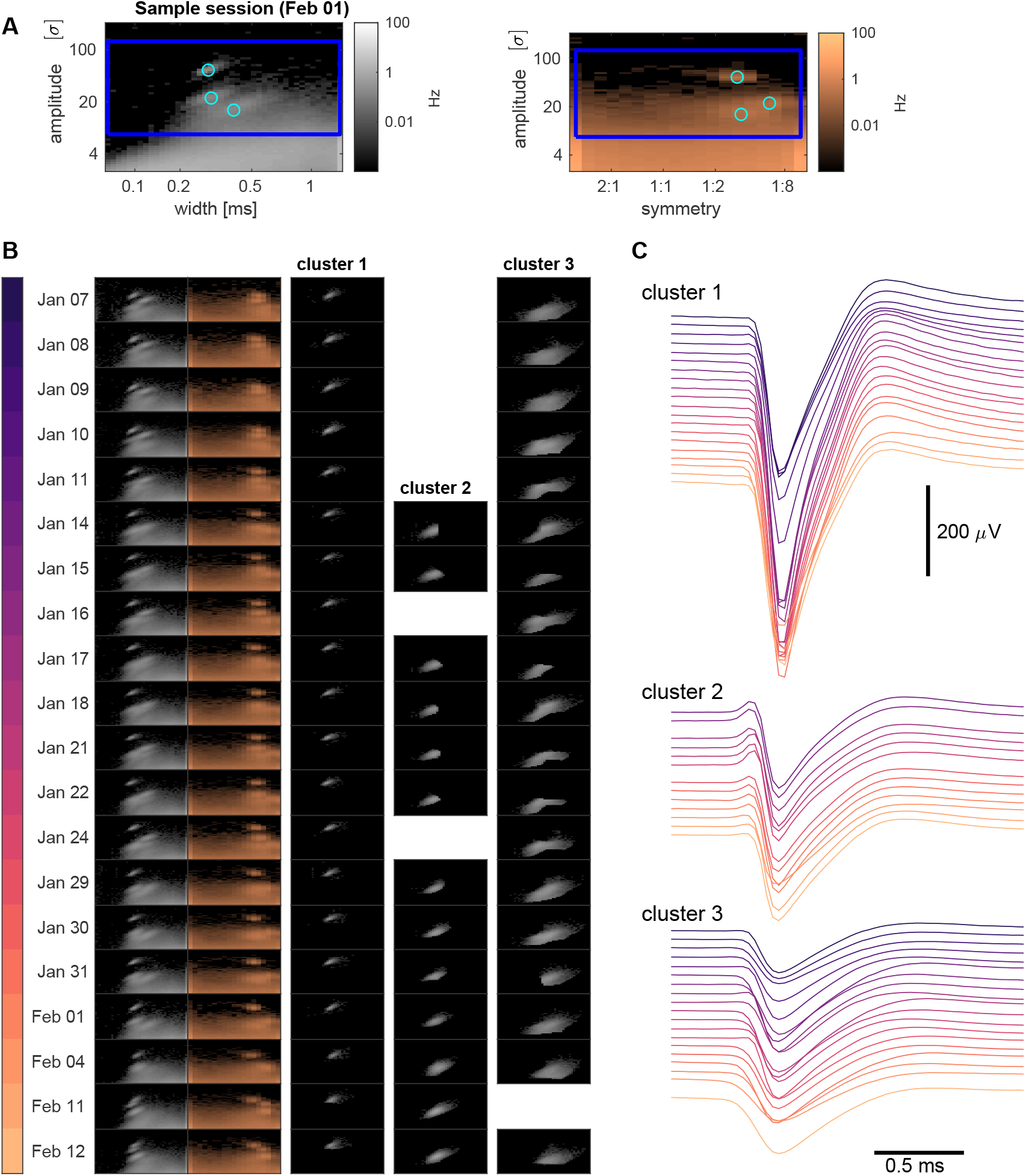
Example of merging clusters across sessions. (**A**) Histograms for amplitudes and widths (left panel) or symmetries right panes) of detected events on February 1. Regions outlined in blue are shown for a range of dates in (B), using the same color code and axes. Cyan circles mark the three clusters detected in this session. (**B**) Left: marginal histograms of local maxima for 20 consecutive recording sessions, labeled with dates. Right: temporal matches of the 3 clusters found on February 1. (**C**) Waterfall plots of average spike shapes, for dates as color-coded in (B). Data from marmoset B. **Figure 2–source data 1**. Source data to generate this Figure

Specifically, if the brain tissue was held in place by the 16 electrode shanks of the array such that relative movements between the electrodes and the sampled neurons rarely happened, we would always record from the same neurons and see identical spike shapes. Otherwise, if there were substantial shifts in relative position between brain and electrodes, both amplitude and spike shape would shift with movement, and we would be unable to track units across a large number of sessions.

We were indeed able to systematically match units across recordings. This was done quantitatively, using the Jensen-Shannon divergence as a distance measure in the histogram shape space (allowing for small amplitude shifts under a penalty). Figure 2 B shows an example of tracking the 3 units observed on February 1 across multiple sessions. Cluster 1 provides an example of a clearly isolated unit with very large spikes with distinctive features, which lasted for about 5 weeks. For this cluster, averaged spike shapes were very similar across recording sessions, with smaller amplitudes for the initial and final recordings (Figure 2 C, cluster 1). Cluster 2 represents a cluster with more modest amplitude spikes and relatively common spike shapes, resulting in somewhat more variable sorting performance. While being reasonably well-isolated from January 29 to February 1, it is contaminated to a variable degree with spikes from different units in other sessions and couldn’t be separated from another cluster in two intermediate recording sessions. Cluster 3 had low spike amplitudes, but would be considered a decent multi-unit cluster from January 29 to February 1. For the other sessions, there is a small local maximum in the shape histograms, but the cluster would be considerably contaminated with unclassified, smaller amplitude spikes. Given that larger amplitude clusters slowly (and independently) drift over time, we can assume that the same happens to units in this cluster, making it difficult to obtain exact matches across recordings. But, the relatively moderate firing rate of the cluster would suggest that few units with defined shapes were involved, distinguishing it from unclassified spikes.

These three example clusters from a brief phase of recording demonstrate both the successes and the challenges of this approach, leaving the real work to be quantifying the overall performance and aligning particular scientific questions with corresponding tradeoffs between unit isolation, data per unit, and number of total units. For example, for the assessment of basic physiological mapping and tuning in cortical areas with known columnar architecture, a mixture of singe units and tuned multi-units is often scientifically acceptable, and this approach could provide a wide array of such units, which is important for thorough functional assays. At the other extreme, questions regarding interneuronal correlations can require confidently isolated single units; this approach would provide a smaller number of units, but a large amount of data per unit (as acquired across sessions), which could provide critical statistical power for these sorts of detailed questions.

In conclusion, our main result is that matching simple shape statistics of spike waveforms across several recording sessions using N-form arrays in marmosets is feasible, and for some units this consecutive recording is possible over notably long periods of time (> 1 month). This grants us the capacity to combine data from multiple experimental days, which we deem “supersessions”. Having demonstrated feasibility, we now turn to the issues of validating and quantifying the performance of this system.

### Tuning properties on individual electrodes are stable across sessions

We further confirmed the stability of the measured “supersession” neuronal activity by evaluating the cross-session consistency of physiological tuning properties. This evaluation was done for the MT array implanted in marmoset J, where we were able to confirm that several sites on the array showed directionally-tuned activity in response to moving dots in the left visual field (as expected when recording from area MT in the right hemisphere).

The MT electrodes recorded strongly tuned multi-unit activity, so we focused on MUA super-sessions for this analysis. We again used our parameterized representation of spike shapes to determine a region of interest (Figure 3 A, E, outlined in black) in spike shape space with strong directional tuning across recording sessions (Figure 3 A, E). This was feasible because tuning on a given electrode was consistent across a wide range of spike shapes (Figure 3 B, F). For the two MUA sites shown as examples, the direction tuning curves measured were stable over almost 3 weeks. This stability of physiological properties, built on top of the stability of spike shapes themselves, further strengthens the case for the validity and viability of supersessions.

**Figure 3.**
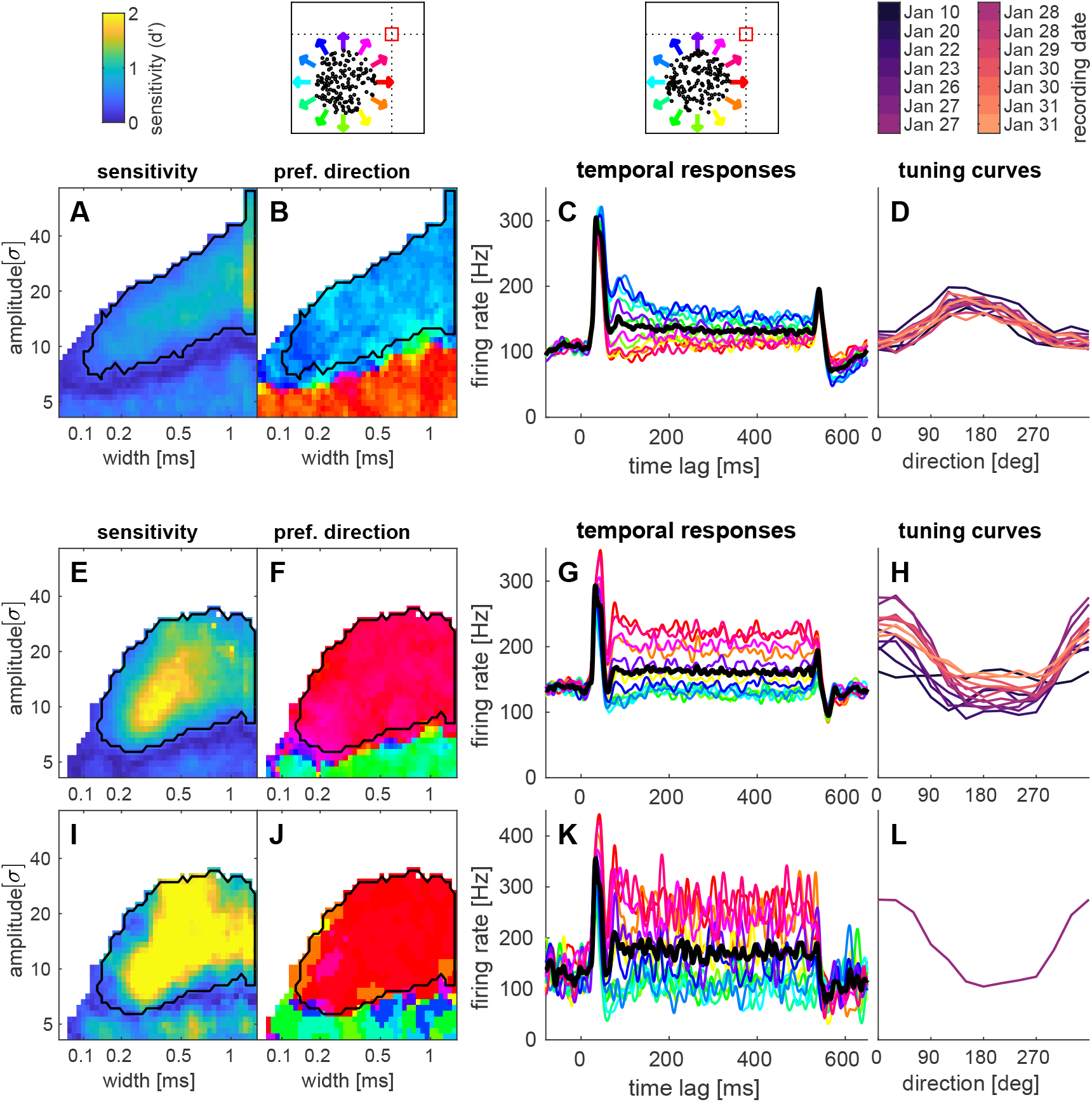
Examples of direction tuning on two electrodes. Top: Legends and stimuli for the examples below. Moving dots were presented at (−15,-15) degrees from the fixation point (red square). (**A**) Sensitivity indices and (**B**) maximum response directions as a function of spike shapes. (across sessions, corrected for a cross session baseline effect). The region outlined in black was used for further analysis. (**C**) Temporal firing rate responses, averaged across sessions and shown for individual tuning directions (colored lines, black line: avg. response, 4041 trials). (**D**) Tuning curves obtained for individual recording sessions (labeled above, some dates had a morning and afternoon session). (**E – H**) Same analysis for a second example electrode. (**I – L**) Tuning observed in a single session (January 27 afternoon session, 254 trials). Recordings in area MT (marmoset J). **Figure 3–source data 1**. Source data to generate this Figure

We therefore created supersessions across these sessions that exhibited stable tuning and spike shapes, which allowed us to combine larger amounts of data for a single analysis. As an example here, we show that supersessions allow us to resolve the detailed time course of responses to individual motion directions at a high temporal resolution (Figure 3 C, G). Note that transient aspects of the motion-driven response were very short and consisted of only a few spikes per trial, such that averages across many trials were beneficial. To illustrate this effect, we show the same analysis for responses obtained in a single session (Figure 3 I-K). Averaging over the temporal responses, we then obtained tuning curves for individual sessions (Figure 3 D, H, L).

In this example, tuning was stable for considerably longer than one week. This demonstrates not only that shape clusters with high amplitudes were stable across sessions, but also that functional properties of low-amplitude activity were conserved across many sessions. Furthermore, being able to combine 10 or more sessions provides an order-of-magnitude increase in trial count that, even assuming some degree of lower-quality unit isolation, should counterweight the relatively short individual behavioral sessions. We delve into this issue in more depth at the end of the results sections.

### Most units in a given recording were observed for several sessions

Having established stability of both spike waveforms and physiological tuning, we now turn to report a more comprehensive statistical description of recording stability and our ability to distinguish spike shape clusters (i.e., to isolate one unit from another). A summary of all tracked units across recording sessions is shown in Figure 4. Spike clusters were regions in 3D-shape-histograms, consisting of a set of voxels, which could be divided into boundary voxels (adjacent to a voxel outside the cluster) and center voxels. If the average spike count in boundary voxels was less than 3/4 of the average density in center voxels, clusters were considered as “better-isolated” and shown in darker colors in Figure 4.

**Figure 4.**
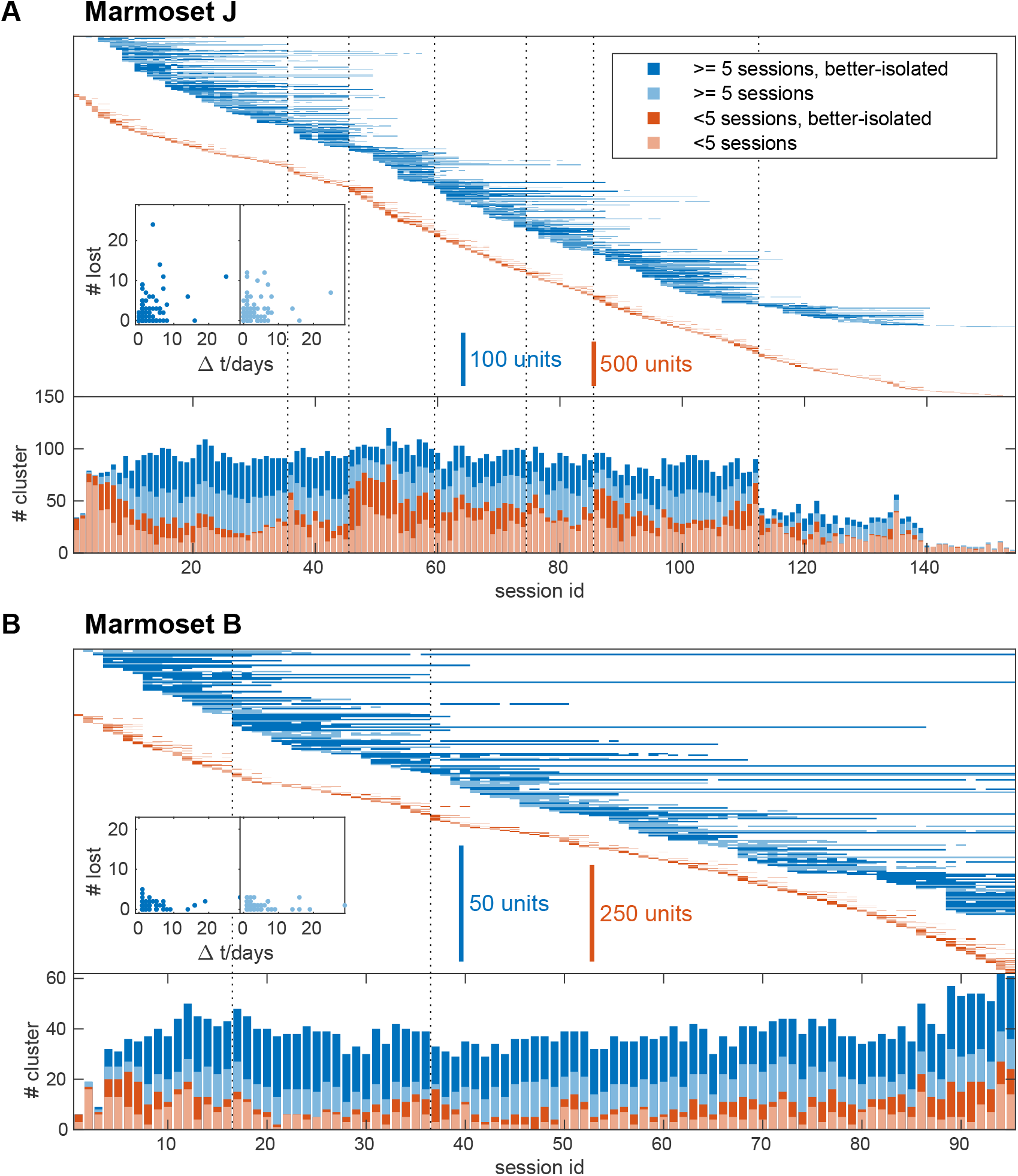
The majority of clusters survives for multiple sessions. (**A**) Clusters detected in recordings of area MT (marmoset J). Top: temporal pattern of long-term (at least 5 sessions, blue) and short lived (*<*5 sessions, orange) clusters. Better-isolated clusters are shown in darker shades. Dotted lines mark times when more than 16 long-term units were lost. Inset: Number of disappearing units as a function of the temporal gap between two recording sessions. Bottom: Number of clusters in each session. (**B**) Same plots for recordings in PPC (marmoset B), except that dotted lines mark times when the highest observed number (five) of long-term units were lost. **Figure 4–Figure supplement 1**. False discovery rate estimates for marmoset J. **Figure 4–Figure supplement 2**. False discovery rate estimates for marmoset B. **Figure 4–Figure supplement 3**. Long-term statistics for marmoset J. **Figure 4–Figure supplement 4**. Long-term statistics for marmoset B. **Figure 4–source data 1**. Source data to generate this Figure and the associated Figure supplements

We further distinguished clusters that lasted for shorter numbers of sessions (*<*5, orange) and longer numbers of sessions (blue, 2.5), as many of the short-lived units had low amplitudes and were less reliably detected.

We found that a large proportion of units in a given recording survived for multiple recording sessions (histograms in Figure 4, blue vs. orange), especially when they were considered as better-isolated (Figure 4, darker colors).

A more detailed visualization of the survival of individual units is shown in the upper half of both panels in Figure 4. This plot can resolve whether the appearance or disappearance of units between two sessions happened locally (i.e., affecting only some individual units), or globally (i.e., affecting most, if not all, units across the array). To further see whether the temporal separation (i.e., number of days) between consecutive sessions was a major factor for the loss (/turnover) of units, we visualized the relation between the number of long lasting units lost and the temporal separation between the two sessions when the loss occurred (Figure 4, insets). Although larger temporal separations tended to correlate with a higher turnover of units, substantial unit turnover could also occur even with very short temporal separations between sessions.

This analysis also highlights a difference between the two animals: while there are several distinct time points of high turnover in marmoset J (Figure 4 A, dotted lines mark disappearances of more than 16 long-term units between consecutive sessions, likely indicative of discrete changes in electrode array position), no such events could be identified in marmoset B (Figure 4 B, dotted lines mark disappearances of the maximum of 5 long-term units, likely indicative of only smaller and/or more gradual changes in array position within the brain). Although we are not sure why the array stability was different in the two animals, this does show that: (a) our analysis scheme is capable of revealing changes and differences in stability; and (b) regardless of whether an array was stable over longer or short terms with or without distinct temporal changes, it is possible to follow units across supersessions in both regimes.

We further quantified how often the algorithm would incorrectly classify two units as being the same, by attempting to merge clusters found on different channels. While such chance matches (Figure 4 – Figure supplements 1 and 2) were unable to explain the number and longevity of units we observed, they did vary considerably across clusters, as some spike shapes were more likely to be found in the data.

Alternatively to asking how well units matched across sessions, we could ask how much long-term units varied over time. Specifically, we were interested in the variability (or coefficient of variation) of properties which were rather neuron and less network specific. Spike shapes or spike amplitudes (Figure 4 – Figure supplements 3 A and 4 A) were used in the process of merging units across sessions and variability would therefore be biased to lower values. Spiking statistics was not used in this process, and we estimated firing rates (Figure 4 – Figure supplements 3 B and 4 B), as they would not be drastically influenced by experimental conditions. As independent measures, we examined spiking statistics at a fast timescale, arguing that intrinsic neuronal dynamics would be more relevant for the dynamics of bursting behavior than the local network activity. We estimated the maximum instantaneous spike rate in a 50 ms temporal window after a spike, relative to the firing rate of a unit (referred to as ‘burstiness’, (Figure 4 – Figure supplements 3 C and 4 C), and the time to reach 75% of this rate, which we refer to as ‘relative refractory period’ (Figure 4 – Figure supplements 3 D and 4 D).

All these measures are expected to fluctuate (due to different behavioral conditions, different levels of recording noise, homeostatic changes in neuronal properties and stochastic errors in the estimates), but would on average be even more different between different neurons. We therefore quantified how much of the variability of these four measures was found across sessions in the same unit, as fraction of the variability across sessions and units (Figure 4 – Figure supplements 3 E and 4 E). While we have no ground truth data for how much variability to expect, we report these numbers here and note that further studies would be required with better constrained marmoset behavior or at least longer recordings in individual sessions, especially for interval statistics at a fast temporal scale. We note that in all cases, most of the variance observed across the population was explained by unit identity.

Figure 5 shows descriptive histograms of the basic properties of all detected shape clusters (grayscale background). We distinguished clusters that survived short-term (upper row) and long-term (lower row). Several basic relations become apparent from visual inspection. First, the spread (avg. diameter) and firing rates of clusters tended to be larger for smaller amplitude waveforms, likely reflecting the effects of merging overlapping shapes from multiple units. Second, large amplitude waveforms were generally more skewed than those with low amplitudes, likely reflecting our descriptive approach’s ability to identify the basic shape of individual unit waveforms. Third, waveforms from the array in MT tended to be narrower than those from the PPC array (two sided Wilcoxon rank sum test, short-term units: p=2e-20, median widths 0.28 ms vs. 0.40 ms and long-term units: p=4e-19 median widths 0.24 ms vs. 0.32 ms), perhaps revealing a biophysical difference that our approach is capable of picking up.

**Figure 5.**
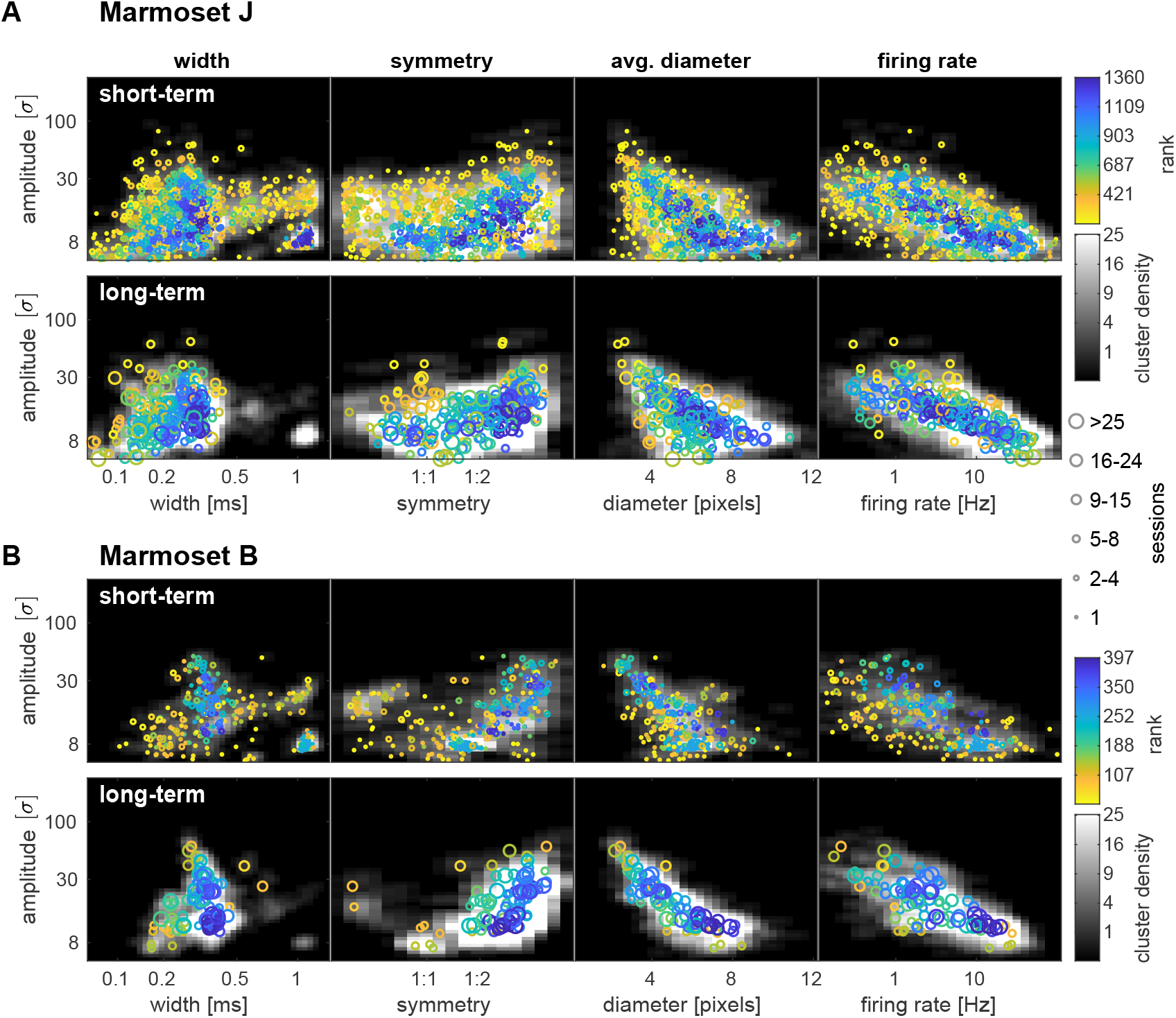
Detected shape clusters are similar (at a population level) when observed for multiple sessions. (**A**) Clusters detected in all recordings and electrodes of area MT (marmoset J). Grayscale represents the density of all detected clusters without merging them across sessions. Colored circles represent individual, better-isolated clusters, merged across sessions. These were ranked according to the corresponding overall density of clusters (i.e. grayscale background) and this ranking is shown in color. Specifically, properties of clusters depicted in yellow were rarely observed and those in blue were commonly found in the data. Clusters surviving less than (top row) and at least (bottom row) 5 sessions are plotted separately for clarity. (**B**) Same analysis for recordings in PPC (marmoset B). **Figure 5–source data 1**. Source data to generate this Figure

Viewing these basic descriptive plots, we also wondered whether long term matches of spike clusters might be a result of detecting different units that just happen to produce similar shapes. To test this, we estimated how likely a given cluster might be mistaken for a different cluster by counting the clusters with similar spike shapes from all recording sessions. We then ranked better-isolated clusters according to the number of similar shaped clusters. The resulting rank a cluster had in the sorted array is depicted in color in Figure 5. A low rank corresponds to isolated units and a low likelihood to detect the same cluster by chance (Figure 5, yellow/green circles), and a high rank means that the corresponding spike shapes were frequently observed (Figure 5, blue circles).

Sorting clusters in this way allows us to investigate whether clusters with commonly observed spike shapes would show a bias in long-term survival. We observed that many clusters with unique shapes survived less than 5 sessions (Figure 5, yellow circles). However, we also noticed that many of these clusters had uncommonly wide or narrow spike widths or very low firing rates. We therefore performed a second ranking, which only included units with an average width between 0.1 – 0.5 ms and an average firing rate above 0.5 Hz and assigned the excluded units the ranks of the next lowest ranked included unit. This was not done to exclude units from our analysis of the relation between spike waveform uniqueness and lifetime, but to group them more evenly.

In order to assess whether clusters with more or less common waveform shapes might show a difference in their lifespans, we analyzed cluster survival, excluding different amounts of the most common cluster shapes. Due to the limited amount of data, we visualized the expected additional lifetime at a given age, assuming a constant probability to lose a cluster in each session. Figure 6 shows that this assumption is reasonable, as the expected lifetime does not change dramatically after 5 sessions. Importantly, except for clusters with the 10% most uncommon shapes, the rate at which spike clusters were lost over time did not depend on how common the spike shapes of that cluster were. This is good news, as it does not appear that the longevity of units over sessions is strongly confounded by the appearance and disappearance of units which happen to have similar spike shapes.

**Figure 6.**
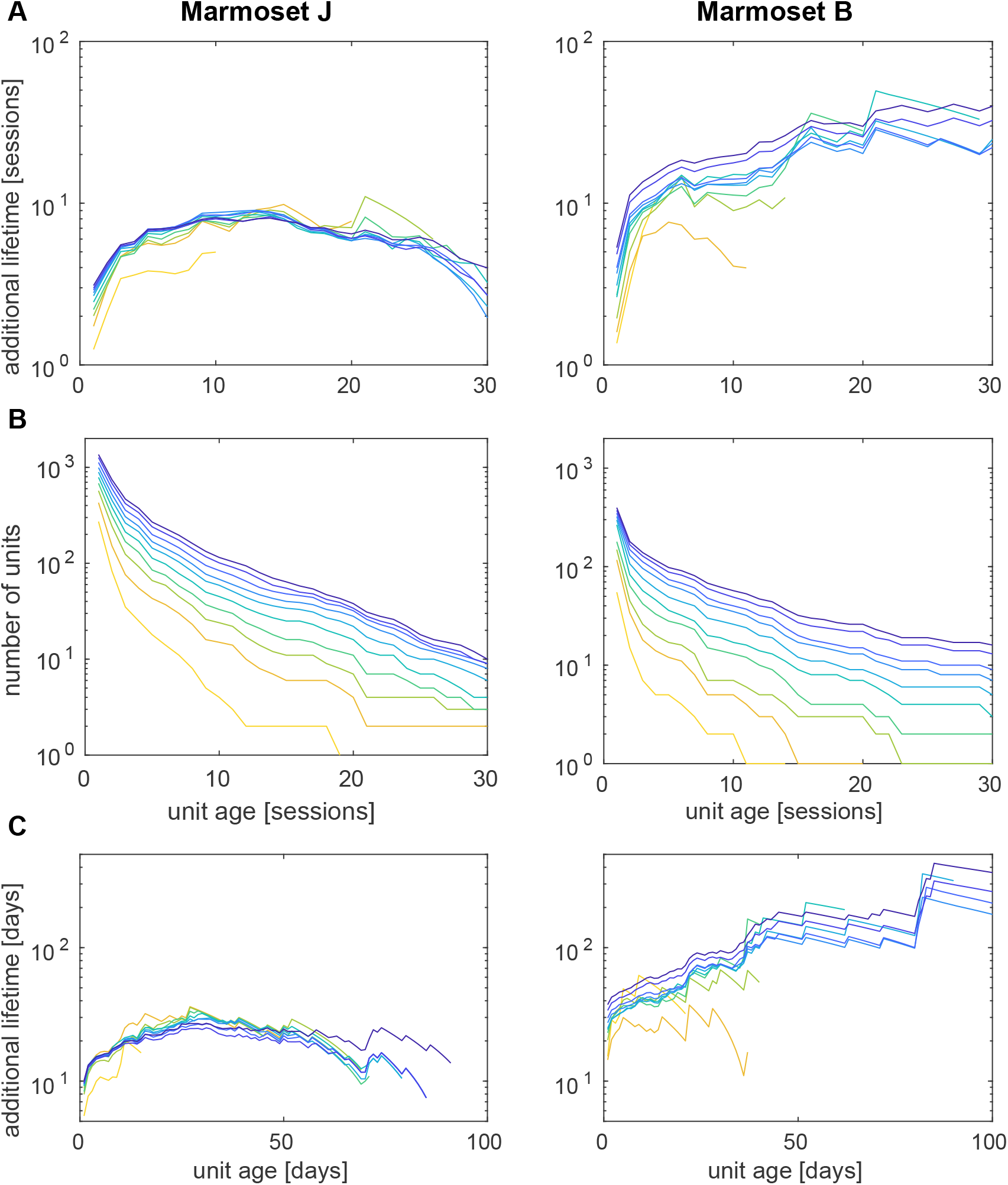
Cluster survival is not an effect of common spike shapes. (**A**) Estimated additional lifetime of clusters after surviving the number of sessions indicated on the x-axis. Coloured lines correspond to the fraction of clusters included in the analysis (steps of 10%, as in Figure 5), where the most yellow curve corresponds to only including the 10% most uncommon shapes. (**B**) Number of units observed for a minimum lifetime. (**C**) Same as in (A) when measured in days rather than sessions. Recordings in area MT (marmoset J, left column) and PPC (marmoset B, right column). **Figure 6–source data 1**. Source data to generate this Figure

This analysis also revealed an interesting difference between the two animals: For the array in PPC, cluster survival was about twice as long as for the array in area MT. Although there were more clusters observed for the MT array, we also observed greater variations in signal amplitude and we gradually lost signal in the later recordings of that array (Figure 1 A). We therefore infer that the observed effect could have been due to a higher degree of general instability of the MT array over time.

### Supersessions provide the power to estimate spatial and temporal aspects of responses across sessions

Finally, we tested whether clearly isolated units could be matched across multiple sessions to assess their spatial and temporal properties. We therefore performed generic receptive field mapping assays at regular intervals over multiple experimental sessions. As proof of concept, here, we describe an example in which both spatial receptive fields and temporal dynamics of responses were estimated using supersession data.

Figure 7 shows two example units. The first unit had well isolated, high amplitude spike shapes (Figure 7 C,E) and a pronounced refractory period (Figure 7 F) for at least 6 recording sessions (firing rate (1.7 ± 0.2) Hz; avg. spike count per trial (400 ms) 0.7 ± 0.4 overall and 1.5 ± 0.5 for stimuli in the receptive field). It consistently responded transiently to stimuli in the left visual field, 50-80 ms after stimulus onset. The second example ((Figure 7 G-L) shows a unit with an amplitude gradually increasing and decreasing across sessions. Corresponding to an increase in SNR and lower contamination by false detections averaged spike shapes became sharper for sessions with large spikes (Figure 7 K). This unit had a much faster response around 40 ms, consisting of about 1 spike per trial (and eventually a slightly elevated sustained activity during stimulus presentation). In both of these cases, the response properties of the unit would have been difficult to determine using only a single session’s worth of data, due to the low absolute number of spikes recorded. For example, the total number of spikes recorded in the first 400 ms in the receptive field of the unit in a single session was just 20-80 spikes, the total number of spikes across all trials about twice that amount. But by evaluating data across sessions, the supersession data shows that these units had clearly-localized receptive fields.

**Figure 7.**
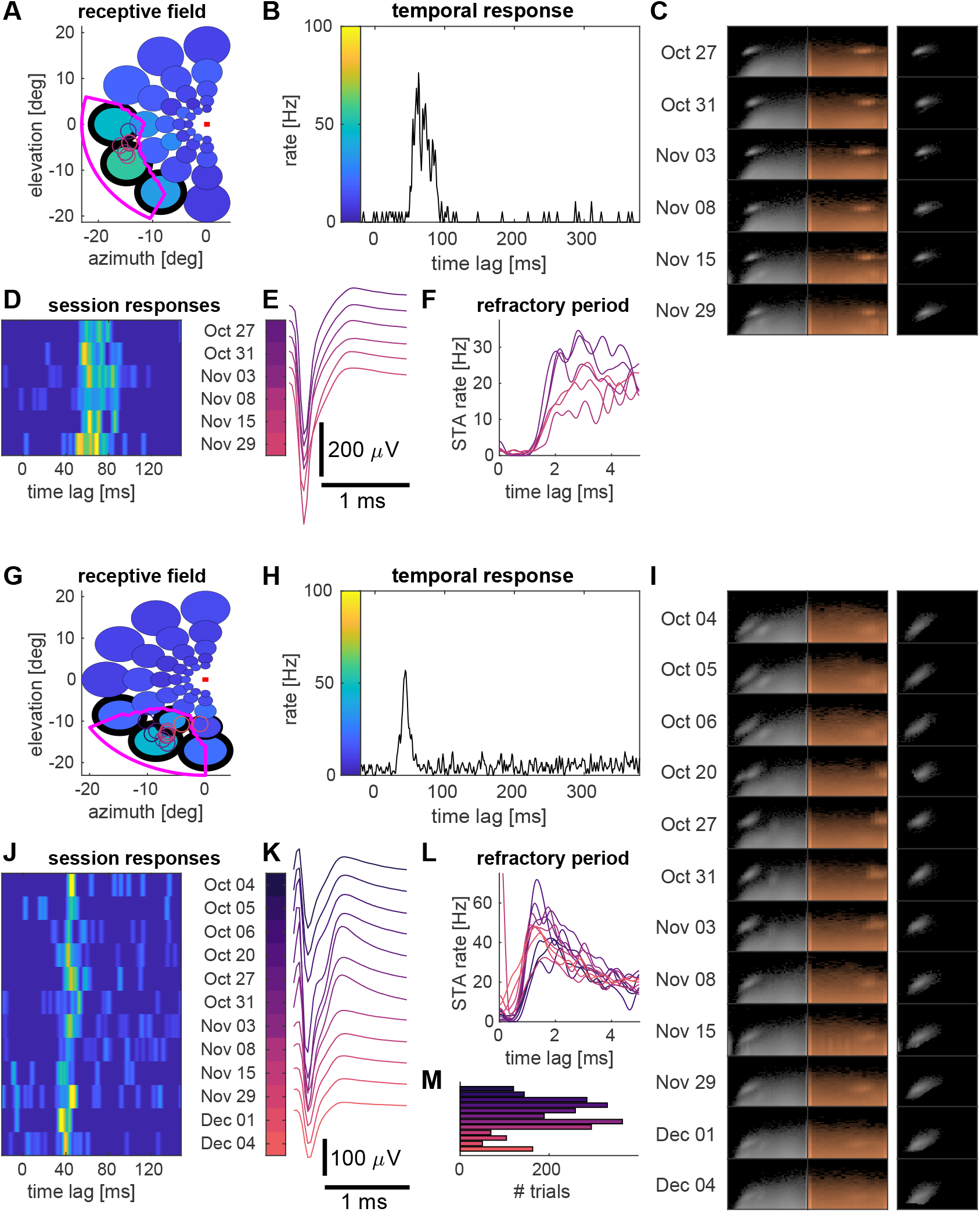
Examples of receptive fields of two units near area MT. (**A**) Maximum firing rates in response to presentation of a disk of moving dots (diameter scaled by 1/2 for clarity; colors indicates firing rate) at a given location in the visual field (fixation spot indicated by a red square). The receptive field (region where the interpolated firing rate exceeded a threshold; see Methods) is outlined in magenta. Colored circles represent estimates of receptive field locations for individual recording sessions. (**B**) Average firing rate for the three conditions (around the RF) outlined in black in (A). (**C**) Marginal shape histograms (as in Figure 2). (**D**) Close-up for firing rates shown in (B) for each recording session. (**E**) Averaged spike shapes. (**F**) Spike triggered averaged firing rates show a refractory period after spikes. (**G-L**) Same as (A-F) for a different unit. (**M**) Total number of trials per session. Colors indicate recording dates (sessions) and firing rates, respectively, and are matched across panels. Recordings near area MT (marmoset J). **Figure 7–Figure supplement 1**. Statistics for aggregate data. **Figure 7–source data 1**. Source data for this Figure and the associated Figure supplement

We further investigated how these examples would generalize to a larger population of units with substantial inhomogeneity in both receptive fields and signal-to noise ratio. For this analysis, we found 172 units that were recorded across at least 4 sessions in which we mapped receptive fields. In order to see whether there was consistency in responses across sessions, we estimated receptive field locations for individual sessions and calculated a ‘sensitivity index’ to quantify the strength of the spatial tuning.

Units that were spatially selective generally had receptive fields that were clustered in a small region of the lower left visual field (Figure 7 – Figure supplement 1 A,B). Importantly, we found that receptive fields were even better localized across sessions in individual units than across the population of equally well or better tuned units (Figure 7 – Figure supplement 1 C). In addition, we saw that the strength of tuning, (quantified as ‘sensitivity index’, see Methods) generally matched between sessions (Figure 7 – Figure supplement 1 D).

While this final analysis outlines a strategy to perform analyses on multi-session and multi-unit data and quantifies consistencies in receptive fields across sessions, we don’t have an obvious reference or gold standard that these numbers could be compared to. These results rather demonstrate what is currently possible, with available data. We do believe that this approach will only improve quantitatively, as array technology continues to improve and yield higher-quality data.

## Discussion

Modern neurophysiological studies in primates require increasingly large amounts of data, either because the parameter space of relevant stimuli or behaviors grows richer (and hence, data are distributed across a larger number of conditions), or because the goal of the experiment itself is to measure more detailed aspects of population activity (and hence, more data are required to estimate higher order statistics). Here, we established the potential of chronically-implanted 3D electrode arrays, coupled with a simple unit identification scheme, to allow for the creation of supersession datasets that transcend the standard limitations of marmoset behavior within individual experimental sessions. We found that high quality activity was evident on this type of array for many months, that a mixture of stable SUA and MUA data could be collected spanning multiple individual sessions, and that these supersessions yielded stable physiological characterizations that were more detailed than those from single sessions.

### Recording performance

With the goal of making the marmoset more strongly viable for detailed quantitative studies, we aimed to develop an analysis pipeline that would be robust to different levels of recording quality, measuring single-unit activity where possible, but at the same time considering multi-unit activity. When applying this analysis to data recorded from implanted electrode arrays over the course of more than 9 months and averaging across all recording sessions, we obtained 28 better-isolated units/array/session. For individual arrays, these averages were 32 and 23 for marmoset J and B, respectively, 20 and 18 of which would be seen across a span of five or more sessions. In addition, we found another 40 and 16 multi-unit clusters per array per session for marmosets J and B, respectively; 18 and 9.5 sessions of these multi-unit clusters lasting for five sessions or more).

In comparison, previous reports of recording stability using planar (2D) ‘Utah’ arrays in macaques (***Dickey et al., 2009***; ***Vaidya et al., 2014***; ***Fraser and Schwartz, 2011***) focused on single unit activity, which strengthened their claims to be able to track individual units, but at the cost of discarding multi-unit activity. Values reported in those prior studies were at most 137 units/array/session, but with large variations across arrays and with decreasing number over time, the average values were closer to 30 units/array/session. In addition, most recordings were done in the first two months after implantation, possibly implying a quicker falloff in signal quality than we encountered with different arrays, and making the comparison to our unit identification and quality less direct.

Although a complete comparison between these types of array is beyond the scope of this proof-of-concept tool introduction, we believe it is likely that the variations in performance observed with ‘Utah’ arrays in macaques were larger than for the 3D arrays we used. In fact, in marmosets, arrays with similar sizes as the ones used in this study (but with fewer electrode contacts) have been reliably implanted and often measured spiking activity for months (***Debnath et al., 2018***).

We conclude this comparison by noting that we recorded from a similar number of units as reported for the larger 96 channel ‘Utah’ arrays (***Dickey et al., 2009***; ***Vaidya et al., 2014***; ***Fraser and Schwartz, 2011***), but from a smaller region of the brain, largely thanks to the denser 3D geometry of the arrays. This is another advantage on the hardware side of this tool, as it allows for larger scale recordings within small brain areas in the marmoset– arrays built for larger primate brains will often sparsely sample within a single area, spanning their footprint over many adjacent areas.

### Long-term stability of units

The 3D array recordings had excellent long-term stability, which is a novel and important result for studies using marmosets. The feasibility of long term recordings is itself not totally unprecedented, as there are multiple approaches that align with our observations in a number of species. Here we review some examples, not just to bolster the case that long term stable recordings can be made in a number of species, but to point to the broader potential adoption of the supersession analysis approach we have introduced.

For example, ***Jackson and Fetz (2007)*** used microwires and studied stability of single units in continuous recordings using a window discriminator, and found single units surviving for up to 17 days in a one year experiment, where microwires were moved periodically to different neurons to improve signal quality. More systematic experiments addressing long-term stability of individual units were done with ‘Utah’ arrays by matching spike waveforms and inter-spike interval histograms across recording sessions (***Dickey et al., 2009***; ***Vaidya et al., 2014***), eventually in combination with correlations and firing rates (***Fraser and Schwartz, 2011***) to increase statistical power. While comprising relatively small numbers of units and recording sessions, these studies demonstrated a few single units being recorded for months, suggesting that there was likely no relative movement between the electrodes and the neural tissue. ***Linderman et al. (2006)*** used continuous recordings to study short-term changes of spike amplitudes and reported moderate amplitude fluctuations in two example units.

The N-form arrays we used had the same spacing between shanks as the ‘Utah’ type of array — albeit with a higher density of recording sites along a shank, and far fewer total shanks. Even though the N-form arrays comprised only 16 shanks, we found a similar long-term stability for well-isolated single units, suggesting that this number of shanks is sufficient to mitigate substantial array drift. The smaller “bed of nails” also permits a slow insertion method, which we hypothesize is important for avoiding damage associated with ballistic insertion methods, especially important in the smaller and more delicate marmoset brain.

In assessing the usefulness of supersession unit data, we used relatively relaxed criteria for unit selection. Given this liberal approach, we did not focus on comparing session-scale average spike waveforms (as these are sensitive to varying amounts of other-spike contamination and noise), but rather distributions of a parametric representation of spikes, where contamination could be considered as a mostly flat, additive component. Likewise, we dropped the comparison of inter-spike interval histograms, firing rates and correlations. While these can provide useful information about unit identity, they rely on a high SNR and good isolation of units in every single session and might even depend on the animal’s engagement in experiments. To avoid discarding large amounts of good data without further inspection, we argue that these measures might best be used for post-hoc tests. Spike shapes themselves proved to be reasonably informative about cluster identity, and for short experimental sessions and low firing rates, multiple sessions may be required to obtain useful second order estimates.

Recent studies in rodents have been very successful in long-term tracking of neuronal activity. However, this performance was in large part made possible by increasing the density of electrode contacts, and therefore the number of observables available for spike sorting. Specifically, ***Okun et al. (2016)*** successfully sorted concatenated data for a small number of sessions and immobile NeuroNexus silicon probes with 4-8 tetrodes (slow insertion). Tetrode recordings in mouse (***Dhawale et al., 2017***) have been used for continuous tracking over weeks. Continuous tracking seems required here due to larger fluctuations in electrical coupling of neurons to electrodes. Recent work with high density arrays (***Chung et al., 2019***) in rats showed smaller fluctuations and allowed sorting segments of data and linking these together. Other recent high-density recording techniques using ultraflexible mesh electronics (***Fu et al., 2016, 2017***) and silicon high-density arrays (***Jun et al., 2017b***) have not yet been systematically studied for unit longevity. In primates, heptodes have been used in acute recordings, in marmoset cerebellum (***Sedaghat-Nejad et al., 2019***) and in macaques ***Kaneko et al. (2007)***, and single unit tracking was done in the latter case.

In terms of stability of units, the following general picture emerges: wires and tetrodes drift within days, but stability is better when they are left in place without an attached micromanipulator ***Okun et al. (2016)*** or when they are continuously tracked (***Dhawale et al., 2017***), approaches which can yield stability for days to weeks. Multiple shanks likely reduce electrode drift and units can be tracked for weeks to months (‘Utah’ arrays potentially for months if no degrading signal quality, ***Vaidya et al. (2014)***; ***Fraser and Schwartz (2011)***), while ultraflexible, polymer based electrodes might remain stable even longer. Our results fit well into this picture.

### Implications for experimental planning and spike sorting methods

Long-term stability offers the potential to generate detailed characterizations of neuronal behavior, but it also requires more careful experimental planning. In the two sections below, we highlight conceptual differences for experimental planning and spike sorting compared to the classical single-session approach.

#### Experimental Planning

While the general long-term stability and the observation of single- and multi-unit activity did support more data-rich analyses than would have been possible from a single session, the fashion in which units ended up being sampled across recordings crucially affects the planning of possible experiments. If, at one extreme, we had recorded from a different set of neurons in every recording session, we would have ended up with a large sample of recorded neurons, but not more data per unit. Such a scenario would allow us to estimate distributions of neuronal behavior in a given area. At the other extreme, if we were to always record from the same set of neurons, we would end up with a small sample, but would be able to measure their responses in many different conditions and further quantify the higher-order statistical interactions between them.

In reality, we found ourselves in a fruitful middle regime: Units were recorded for variable durations, in which a small fraction of units both appeared and was lost between recording sessions. This process was not entirely random, as we saw that most units disappeared during the initial sessions after their appearance. This means that the chance for a unit to survive for another session increased with the number of sessions that this neuron had already been observed. Hence, if we were to ask which of the units we would most likely observe in a future session, the best bet would be those units that were already observed for the most sessions in the past.

The variable lifetimes of units also provide an additional tool for raising the standard for isolation. Restricting an analysis to only long-lasting units would likely reduce the chance of including less clearly isolated units. Such units may not be found in some of the recordings due to variations in signal amplitude.

The exact timescales at which units were lost between sessions varied slightly across our two test arrays/animals. However, there may be two different mechanisms involved: while we found a relatively low, constant turnover of units on both arrays, in marmoset J we additionally saw a few events where a large fraction of units was lost between subsequent recordings (Figure 4). These events could not be explained by a long temporal gap between the recordings, suggesting a relatively fast mechanism for that, with a timescale of hours to days (as opposed to weeks and months). We believe that these findings can impact the planning of experiments using chronic arrays.

In the classical single session approach, experimenters devote part of the experimental time for general characterization of receptive fields and tuning of neurons, in order to target a neuron and adapt the stimulus properties to efficiently sample responses, avoiding stimuli without an expected effect on the neuron’s firing behavior. In the case of chronic array recordings, we record from many neurons with potentially different receptive fields and tuning properties, suggesting the use of more general stimuli, e.g. sampling a larger visual area and different tuning directions. Especially when studying interactions between a small number of units, one should keep in mind that some of these units may disappear during the course of an experiment and it would be advisable to start with a larger group of candidate units. In this regard, chronic arrays would be ideally suited for continuous tasks and naturalistic stimuli (e.g. ***Huk et al. (2018)***; ***Knöll et al. (2018)***), which efficiently sample a large parameter space, allowing for simultaneous characterization of units with different tuning properties.

If, however, an experimental design requires finding persistent units in order to adapt focused studies to suit their tuning, we recommend choosing units that have already been observed for at least 3 sessions, as these units have a high chance to survive the next sessions. In our experiments, such units had a conditional (additional) lifespan of 6 and 14 sessions (for marmoset J and B, respectively, cf. Figure 6 A). Likewise, studies of changes in firing behaviour of single units across sessions (e.g. while an animal is learning a task, or after drug treatment) are in principle feasible. However, such experiments can usually not be repeated in the same animal, and few units will be clearly isolatable, resulting in a rather inefficient use of the acquired data. In this case, the suggested approach is to perform several consecutive studies on an animal, which is possible given the longevity of the arrays used here.

Importantly, we have shown that it is feasible to combine data across multiple sessions to infer tuning properties of neurons from multiple sessions. When looking at a population of recorded units, we would encounter a relatively high variability in both signal-to-noise ratio and physiological properties across the population. Such variations would generally result in different requirements on the amount of data needed for statistical tests (e.g. a weak tuning requires more data to determine a receptive field). It was therefore useful to sort units according to their tuning strength, and to perform a relatively focused analysis to specifically detect changes in receptive field locations with high statistical power, using data from single sessions. This strategy would then allow to ask the more detailed questions for data pooled across sessions in a second step.

The same type of analysis should be possible for inter-neuronal correlations. Our results also highlight that, in many cases, it would be incorrect to assume that units with similar spike shapes recorded on the same electrode in subsequent sessions would correspond to different neurons.

We conclude that chronically implanted electrode arrays allow for both sampling of a large set of neurons and detailed analysis of a few long-term units, but different timescales need to be considered when planning experiments. If the objective is to sample the population of neurons across a brain area, experimental sessions could be separated by a month to take advantage of appearance and disappearance of neurons on the array. If instead the objective is a detailed analysis of a smaller set of neurons and their interactions, daily recordings for 2-4 weeks are ideal.

#### Features of the spike sorting method

We adopted a modular strategy for spike sorting, where individual sessions were processed independently and could be iteratively merged to form ‘supersessions’. In this way, experimenters can perform preanalyses as data are generated and determine receptive fields and tuning properties of neurons to guide stimulus selection as well as monitor recording quality. This modular approach further facilitates excluding particularly noisy segments in individual sessions, which might impair or bias the clustering algorithm.

The primary reason for eschewing existing spike sorting methods was a general concern about robustness when stationarity assumptions were not met across recording sessions. This is a known challenge to even cutting-edge algorithms (***Jun et al., 2017a***). We instead chose a simple parametric representation that was designed to be robust to noise and artifacts, which can differ from session to session. Our focus was on characterizing the peak of the depolarization phase using unimodal templates where the SNR would be highest. While spike shapes can be strongly bimodal, depending on the relative position of the electrode and neuron, the shapes for spikes with highest amplitudes near the soma have been shown to be largely unimodal in theoretical studies (***Lindén et al., 2011***; ***Quian Quiroga, 2009***; ***Camuñas-Mesa and Quiroga, 2013***). As we recorded spikes on single electrodes and could expect a large number of neurons in the vicinity of an electrode (***Pedreira et al., 2012***), high amplitude spikes would be easiest to separate from other units. This situation would certainly be different for high-density probes. The process of estimating parameters of the spike shapes was essentially an optimization. We would shift a template temporally at sub-sampling resolution and change its width and symmetry to best match a local minimum in the raw voltage traces. In practice, this step was implemented by running the raw data through a large filter bank on a GPU.

Our spike sorting approach did not solve the problem of overlapping spikes. However, it greatly reduced the problem as the time interval needed for detection was reduced to the width of the spike and thus, due to zero padding, much smaller than the the width of the templates in the filter bank. In addition, for cases where overlapping spikes exist, we should see them in the shape histograms as somewhat isolated shapes that are a bit wider and of higher amplitude than an adjacent cluster. In our data, we did not find evidence for significant numbers of overlapping spikes near isolated clusters. Overlapping spikes would generally lead to wider and larger observed spike shapes, and such shapes would be reflected as asymmetries in the histograms, where larger and wider than average spikes would be found with a low probability. We didn’t observe such asymmetries, so we can conclude that overlapping spikes were small enough that they wouldn’t affect the observed spike shapes to a greater extent than noise. This situation was different for low amplitude events which could not be separated into distinct clusters, but clearly showed stimulus dependent modulations (as in Figure 3 C, G). These events would necessarily overlap in many cases, as their baseline rate was in the order of 100 Hz and peak rates in single trials therefore likely an order of magnitude higher. Hence, firing rate estimates for low amplitude spikes should be read as a lower bound, providing useful (slightly distorted) information about tuning in sustained responses, while truncating transient responses.

In this work, we used the parametric representation of local mimina as a spike sorting method. But we could certainly perform spike sorting with an existing method and obtain these parametric representations for spikes in order to subsequently match spike clusters across recording sessions. Likewise, as current sorting techniques are validated with respect to stability over long time frames, it would be straightforward to replace our sorting approach. However, our sorting approach could still be used for fast, online assessments of recording quality, neuronal yield and tuning properties as it does not require manual curation.

### Application to data

In many cases, we observed that shape clusters appeared and disappeared gradually over time, such that the observed spike amplitudes were highest around the middle of their lifetime. We could thus have a situation where some shape clusters of a given unit were clearly isolated single unit activity, and others were contaminated (e.g. Figure 7 I). Although this effect means that some of the unit data from ‘supersessions’ is less well-isolated than conventional singe-session data, the framework can also be used to estimate the impact of contamination for a given analysis, and hence to determine in a principled manner how high an isolation standard is required.

To give an example how such analysis could look, assume that we have a number of sessions (W) where a unit was well-isolated, and some sessions (C), where the same unit was contaminated with low amplitude spikes from other neurons and some of its spikes were lost due to low amplitudes. We would then pool data from each group (W and C) of sessions to obtain a larger sample size and estimate firing rates and interspike interval histograms.

Assuming that low amplitude spikes from other neurons are uncorrelated (alternatively, the interspike interval distribution of low amplitude spikes could be estimated with sufficient data) and uniformly distributed, we would fit the ISI histograms of group C as a linear combination of the ISI histogram of group W and a uniform distribution. The component explained by the uniform distribution could then be translated into an estimate of the spike count for the low amplitude spikes from other neurons (i.e., dividing the rate of the uniform component by spike count of group C and multiply with the total recording duration of group C). To obtain an estimate of the number of spikes missed in group C due to low spike amplitudes, one can multiply the difference in firing rates between group W and C with the total recording duration of group C and add the spike count for the low amplitude spikes determined above. After doing a given analysis separately for groups W and C, one could then compare the results and see how they are affected for a known contamination and signal loss.

Furthermore, if one looked into the datasets of group W, one would likely find spikes that are statistically similar to the contaminating spikes in group C, simply by identifying identically shaped spikes at much lower amplitudes. Therefore, it is possible to create surrogate datasets with known contamination (and, by removing spikes, signal loss) and treat them as a model to predict effects on a given analysis. The above analysis would then provide independent data to test this model.

Apart from spike clusters, our sorting approach also gives access to low amplitude spikes that do show tuned responses to visual stimulation, but likely arise from a multitude of units with a continuum of corresponding spike shapes (e.g. Figure 3). For the purpose of decoding neural activity, such low amplitude spikes can be of great value. In fact, results from other groups indicate that lowering the detection threshold increased the performance of a decoder despite losing information about the neuronal identity (***Trautmann et al., 2019***; ***Kloosterman et al., 2013***; ***Todorova et al., 2014***). Our work suggests that we can define a detection threshold (or region of interest) post-hoc, based on responsiveness to stimuli known to drive neural activity. We refer to this activity as multi-unit hash (MUH), creating a third category alongside with MUA, which should form clusters that are separable from MUH, and SUA which would additionally show a clear refractory period. We need to stress here that MUH is still distinct from the ‘unsorted spikes’ often left behind by most sorting algorithms.

In summary, we were able to create ‘supersessions’ for individual units on a timescale of several days to a few weeks. This allows for more statistical power than a single session’s worth of data can provide, and hence could put the awake marmoset preparation more on par with that of macaques. This is important because the marmoset is also a “pivot species” to richer and more powerful techniques that are more difficult to apply to the macaque. Such supersessions do require reconsidering the design of experiments to handle the comings-and-goings of identified units. Such experiments will likely have a long term structure where basic characterization of neural response properties is performed approximately once a week, with the remainder of experimental data collection being dedicated to more sophisticated experiments.

## Methods and Materials

### Electrophysiology preparation

Two marmosets were implanted with N-Form arrays (Modular Bionics, Berkeley, CA, USA) in area MT (marmoset J) or PPC (marmoset B). Prior to placing the chronically implanted array, we drilled a grid of 9 burr-holes over and surrounding the desired brain area based on stereotaxic coordinates from ***Paxinos et al. (2012)***. We performed extracellular recordings using single tungsten electrodes in each burr-hole to fine tune the placement of the array based on the physiological response. The MT array was placed based on high response to direction of motion, while the LIP array was placed based on high eye-movement related activity. A small craniotomy and duratomy were made surrounding the desired area for array placement.

The N-form array was mounted on a stereotax arm and manually lowered till tips of the shanks had entered the brain. The brain dimpled slightly, then the tissue relaxed around the implant. The array was then slowly lowered until the baseplate was just above the brain’s surface. The array was stabilized and sealed with KwikCast before being closed entirely with dental cement and acrylic. The array connectors were enclosed in a custom 3D-printed box embedded in the acrylic implant.

Animal procedures described in this study were approved by the UT Austin Institutional Care and Use Committee (IACUC, Protocol AUP-2017-00170). All of the animals were handled in strict accordance with this protocol.

The N-form arrays (Modular Bionics, Berkeley, CA, USA) consisted of a 4×4 grid of electrode shanks, spaced by 400 µm. Each shank was 1.5 mm long and had 4 electrode contacts, one at its tip, and three more at 250 µm, 375 µm and 500 µm distance from the tip. Extracellular signals were recorded at all 64 electrode contacts with sampling rate of 30 kHz, using the OpenEphys recording system (***Siegle et al., 2017***). For marmoset J, seven of the electrode contacts were found damaged after the surgery and ignored for further analyses.

### Visual tasks and stimuli

All stimuli were presented using custom MATLAB (Mathworks) code with the Psychophysics Toolbox (***Brainard, 1997***) and a Datapixx I/O box (Vpixx) for precise temporal registration of stimulus, behavioral, and electrophysiological events (***Eastman and Huk, 2012***).

Marmosets were trained to fixate a central dot in the presence of peripheral visual stimuli. The animals fixated the dot within a window of 1.5 degree radius for the whole trial to obtain liquid reward in the form of marshmallow juice. If the marmoset broke fixation, the trial was aborted. Fixation was acquired and held for 200 ms before a stimulus appeared.

To measure MT receptive fields, we presented a circular cloud of randomly moving dots for 350 ms at one of 35 different screen locations during controlled fixation. The diameter of the stimulus aperture scaled with the eccentricity of its center.

To measure direction tuning, we presented coherent motion in 12 possible directions at a fixed location based on previously measured receptive fields. Each trial contained motion in one direction for a duration of 500 ms.

For PPC recordings, marmosets were trained to perform a memory guided saccade task. The animals fixated the central dot while a target dot was briefly flashed at a random location in the periphery. After a delay of 400-1000 ms, the central dot was extinguished and the marmosets received liquid reward for saccades to the remembered location of the target. Memory guided saccades are well known to generate PPC activity in primates (***Andersen et al., 1990***). The task itself was not part of the investigations in this work. We outline it here as context for the behavioral engagement of the animal in the experiments and to emphasize its potential to drive neuronal activity in PPC.

On average, recording durations of individual sessions were (26 ± 13) min for marmoset J and (41 ± 12) min for marmoset B.

#### Pre-processing

We filtered a 60 Hz component out of the raw data for each electrode using a custom made algorithm. We also performed common average referencing by subtracting (projections onto) the median of high-pass filtered signals over all electrodes from each channel. We further up-sampled data to 60 kHz before feeding into Kilosort (***Pachitariu et al., 2016***). For this, values between samples were obtained by linear interpolation and values at samples were smoothed with a [1/6 2/3 1/6] smoothing kernel to obtain a uniform variance across data points for the case of Gaussian white noise.

### Spike sorting

Code for the spike sorting pipeline is available at https://github.com/HukLab/SuperSessioning and will further be made available within the SpikeInterface project (https://github.com/SpikeInterface, ***Buccino et al. (2020)***).

We aimed at jointly sorting spike data from tens of recording sessions (marmoset J: N=154, marmoset B: N=95) under the following constraints:

1. Marmosets were head-fixed, but able to move their bodies within the chair, creating temporally variable amounts of noise in the data.
2. Electrodes were separated by at least 2.125 µm and spikes were not generally expected to be seen on multiple electrodes.
3. We observed only few separable units (0-3) per electrode.
4. There was no apparent electrode drift within recording sessions.
5. Spike clusters needed to be matched across recordings.

If spike shapes are known, then template matching would be the best way to detect spikes. However, if spikes are to be sorted, information in the raw data needs to be used to separate spike clusters, and especially to separate them from fluctuations in the background noise level and low-amplitude events of neuronal origin. A good sorting algorithm therefore needs to make estimates that are maximally invariant when subjected to noise. Potential issues are:

1. Baseline estimate: errors could change the match of bimodal templates. This may especially become a problem when the noise level is temporally varied.
2. Sampling frequency and temporal resolution for peak detection: Misaligned spikes differ in shape. This can be resolved by upsampling the data, but results in longer templates.
3. Temporally overlapping spikes: Need to be detected and fitted.

To address these three issues, we generated a bank of unimodal templates (essentially triangles with a tip rounded off by a cosine function) which varied in phase (to effectively yield 180 kHz sampling frequency), width and symmetry (see examples in Figure 8 B), covering a wide range of possible shapes. Each template was normalized to have an energy (sum of squared entries) of one. Using this bank of templates in a template matching strategy reduces baseline errors, temporal misalignment and the chance of fitting overlapping spikes, but does sacrifice some detection power (when compared to using templates generated from the data, about 10% of the signal power).

**Figure 8.**
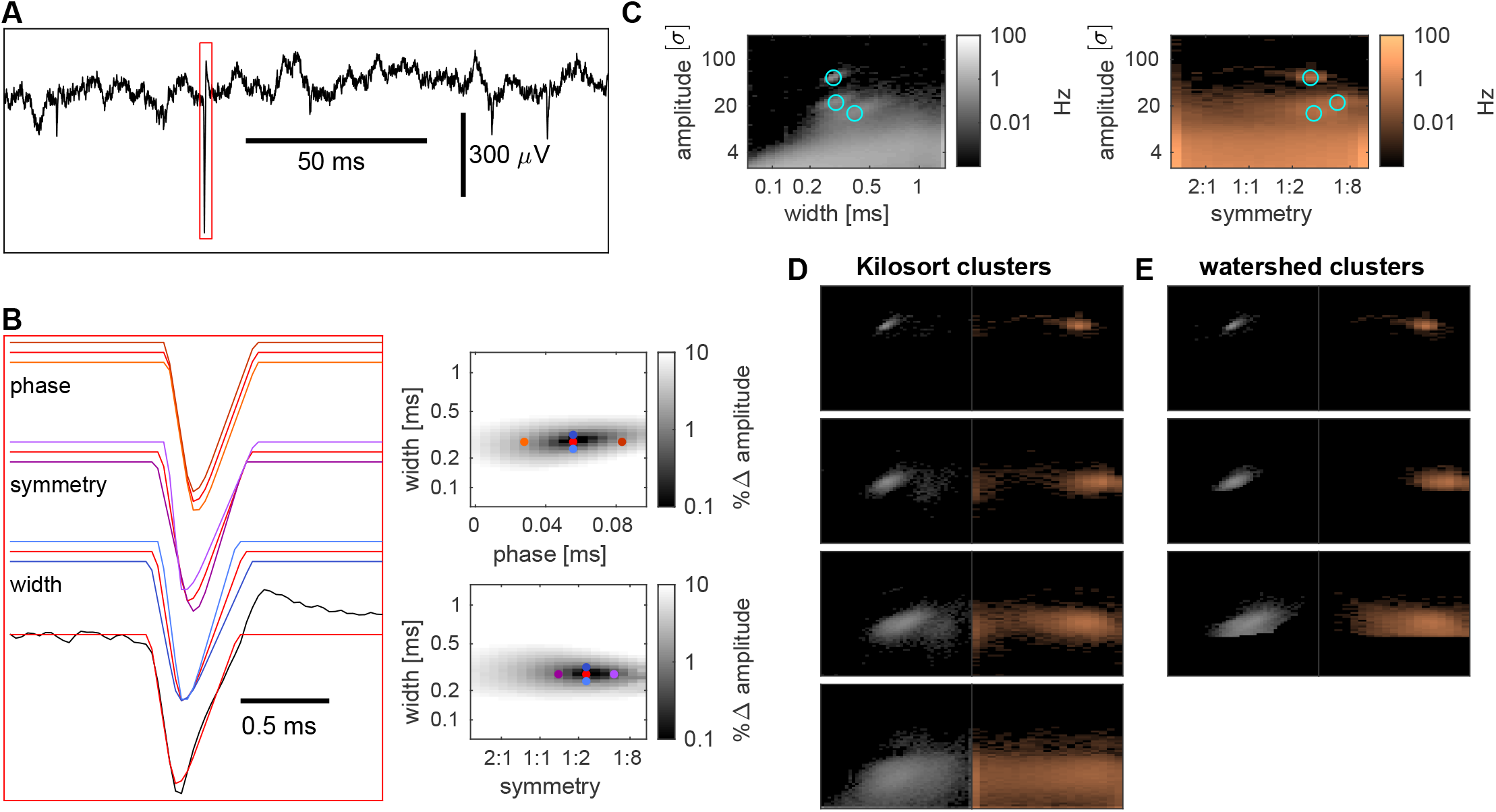
Spike detection and sorting. Raw voltage traces from single electrodes (**A**) are matched in a sliding window to a set of triangular, unimodal templates (examples in **B**, upper left) differing in width, symmetry and phase offset. Local maxima of template - raw trace matches in this parameter space (right plots, dots colored as in left panel) are then detected as putative spikes with a shape characterized by the corresponding width, symmetry and signal power (dot product of template and raw trace). **(C**) Histograms of shapes for an example electrode and recording (marginal distributions). Locations of clusters determined by a watershed algorithm are marked with cyan circles. (**D)** Shapes of events detected by Kilosort on the same electrode, grouped into clusters by an automated procedure. (**E**) Clusters determined by the watershed algorithm (corresponding to the cyan circles in (C)). **Figure 8–source data 1**. Source data to generate this Figure

We determined local maxima (in time and width, but global in symmetry to avoid double detections) for the match (dot product) between our templates and the preprocessed voltage traces. In this setting, we were fitting the peak of the depolarization phase of a spike. While an error in the baseline estimate would have an effect on the detected spike power, it would have little effect on both the estimated spike width and symmetry. Temporally overlapping spikes were less likely as the temporal interval for detection was restricted the duration of the depolarization phase (i.e. 0.5 ms or less) and a linear combination fitting was not necessary in our recordings. Note that we did not capture the repolarization phase of a spike at all, however, we argue that due to smoothness constraints, the shape of the repolarization phase covaried with its symmetry, and its duration was hard to estimate due to potential drifts in baseline. Matching a large set of potential templates was computationally expensive, but also well suited to run on a GPU. Our implementation ran about twice as long as recording the data for 64 electrodes sampled at 30 kHz. Marginal histograms of shapes obtained for an example recording are shown in Figure 8 C.

Clusters of spike shapes were then determined with a density based approach, using the watershed algorithm, which required some amount of smoothing and a step to reduce global density gradients. In more detail, we aimed constraining the number of spikes to average, rather than setting a fixed kernel size for smoothing. For a given number of spikes, we could then estimate the radius required to find that number of spikes, and the watershed algorithm would yield clusters. This approach tends to fail when there are global gradients in spike density and further use extremely small volumes for high spike densities. Therefore, we instead determined the area of a number of spikes that scaled sublinearly with a local firing rate baseline *R*. This baseline was estimated by smoothing with a trivariate Hanning kernel (width 13 bins, truncated first and last bin, and sheared a bit, by 0.5 bins in amplitude per bin in width, such that larger spike widths would be combined with lower amplitudes, to reduce a potential bias due to spike clusters, which were often tilted in the opposite direction). We applied a sublinear scaling and added a small offset to that baseline to determine a firing rate (and therefore the number of spikes), given by 0.015 Hz +0.7*R*^0.9^ for which we determined the required radius. We excluded areas from the analysis for which that orting. Raw voltage traces from single electrodes (**A**) are matched in a sliding window to a set of triangular, n **B**, upper left) differing in width, symmetry and phase offset. Local maxima of template - raw trace matches in, dots colored as in left panel) are then detected as putative spikes with a shape characterized by the and signal power (dot product of template and raw trace). (**C**) Histograms of shapes for an example electrode tions). Locations of clusters determined by a watershed algorithm are marked with cyan circles. (**D**) Shapes of he same electrode, grouped into clusters by an automated procedure. (**E**) Clusters determined by the watershed cyan circles in (C)). data to generate this Figure radius was larger than 5 bins. To avoid instances where the watershed algorithm would turn individual voxels into clusters, we determined a sliding median across 3×3×3 voxels. We further note that there is a dependency between the recording duration and the resolution of this method (i.e. higher resolution for longer recordings).

For clusters obtained from the watershed algorithm (using a three-dimensional 18-connected neighborhood), we excluded clusters that systematically had extreme values for spike width or symmetry or very low amplitudes. Specifically, we ensured that clusters had their center at least half a standard deviation above the lowest or below the highest bin. As there were many events with wide shapes, we lowered the exclusion threshold for wide spikes to half a standard deviation below the second highest bin. For amplitudes we included clusters with and amplitude of at least half a standard deviation above 2.7*σ* for lowest spike widths and 5.9*σ* for highest widths (linear cutoff in the histograms).

To show that these spike clusters indeed corresponded to units found in a conventional spike sorting approach, we sorted spikes with a widely used spike sorting algorithm (Kilosort, ***Pachitariu et al. (2016)***). For that, we used a low threshold for splitting clusters in the Kilosort algorithm and extracted the shapes of the corresponding spikes from our template matching strategy. This allowed us to perform the manual step of merging clusters in an automated procedure, using the Jensen-Shannon divergence between shape histograms as a distance metric.

We obtained three dimensional histograms of shape parameters for spikes from each Kilosort cluster (Figure 8 D). We compared Kilosort clusters to clusters obtained by running the watershed algorithm on shape histograms and found a good match for high amplitude clusters (Figure 8 E). The latter clusters were (by construction) better localized in our histograms and we decided to use them instead of Kilosort clusters in the following analyses.

#### Possible extensions

We implemented the spike sorting for the case of single, isolated electrodes. An extension to dense arrays is beyond the scope of this article, but we will briefly discuss potential implementation issues here.

1. Linear arrays/stereotrodes: can be treated as another dimension, like the phase. This just requires one to set a spatial extent of spikes, creating spatially shifted templates. With this method, one could determine maxima at each time frame for each spatial shift, and do a recursive maximization in a second step to obtain spatially isolated maxima.
2. Spatial grids: memory constraints on the GPU will currently require chunking the array into rows of electrodes.

Our current implementation does not include a template generation and matching step, potentially resulting in suboptimal detection performance. A potential improvement, while still avoiding the baseline issue, could be to generate templates, smooth them with a kernel and generate template versions with different widths and phases by interpolation. We would need to normalize the templates to unit power and reduce positive (repolarization) parts of the templates (e.g. divide by 2), to reduce a potential baseline effect. Then we would replace the predefined templates of a given cluster (obtained from the watershed algorithm) with these templates, while keeping the other predefined templates as alternative options (for events that do not match a particular template). Next, we could rerun the detection with the modified set of templates, considering events which are best matching the inserted templates as spikes.

### Cross-session merges

We computed pairwise Jensen-Shannon divergences between existing clusters from the previous 2 sessions and clusters from the current session allowing for small shifts in amplitude, width and symmetry for a penalty. Specifically, we did multiply the Jensen-Shannon divergence with the inverse of Hanning kernels with a half-width of 7 (for amplitude) and 3 (width and symmetry) bins. Each cluster from the current session was then merged with the existing cluster with the smallest Jensen-Shannon divergence if this was below a threshold of 0.3 ln(2), otherwise it was labeled as a new cluster. To allow for slow temporal drifts, the merged cluster was then assigned a shape density equal to the average of the previous and current density (resulting in effective down-weighting of earlier densities).

### Motion direction tuning

Tuning of spiking activity to the motion direction of a visual stimulus was examined as a function of the width and amplitude of spike shapes, rather than for well isolated clusters, to systematically investigate how much of the low amplitude events was affected by visual stimulation. To this aim, we marginalized over the symmetry parameter of spike shapes, and used a sliding window of 5×5 pixels for amplitudes and widths, to obtain samples of spikes around each spike width and amplitude. For a temporal window from 20-470 ms from stimulus onset, we computed mean and standard deviation of spike counts for trials from each stimulus condition, excluding the 3 highest and lowest spike counts from the analysis for robustness of the estimate. The difference between opposing motion directions in the stimulus was then divided by the root mean squared standard deviations to obtain a sensitivity index for each direction. We maximized the sensitivity index across motion directions and, for a sample session, visualized the argument of the maximum as tuned direction in Figure 3 J and the maximum value as sensitivity index in Figure 3 I. To average these sensitivity indices and directions across sessions, we treated the tuning in each session as vectors in the tuned direction with a length equal to the sensitivity index, and averaged them, to obtain an interpolated tuning direction and averaged sensitivity index, shown in Figures 3 B, F and A, E, respectively. To obtain a region of interest for analysis of all stimulus dependent events found on a given electrode, we thresholded the averaged sensitivity indices at 0.3 and determined connected regions exceeding this threshold. The largest connected region was then used as a region of interest (outlined in Figures 3 A,B,E,F,I,J) for the cross session analysis performed in Figure 3 C, D, G, H, as well as the single session spike time histograms and tuning curves in Figure 3 K, L. All spikes within that region of interest were used to compute spike time histograms with a bin width of 1 ms and temporally smoothed with an 20 ms wide Hanning kernel (Figure 3 C,G,K).

To see how tuning responses at a given electrode site change across sessions, we determined tuning curves for each session (Figure 3 D,H,L). Theoretically, a drift in firing rate or sensitivity could signal a change in coupling between neurons and the electrode, eventually caused by z-drift. Likewise, due to the spatial organization of area MT, a change in phase could reflect a lateral movement of the electrode.

### Cluster survival

Spike shapes were very similar for a large fraction of clusters. It could be that clusters only appeared to last across sessions, but in fact represented multiple different clusters that just happened to have matching shapes. Therefore we wanted to test for a bias in longevity for units with common spike shapes. We computed histograms of amplitudes, widths, symmetry and volume of shape clusters, and the average of these quantities for each better-isolated unit across sessions. We then ranked units according to the local density of shape clusters. A lot of short-lived units had uncommonly wide or narrow spike widths or very low firing rates. We therefore performed a ranking, which only included units with an average width between 0.1 – 0.5 ms and an average firing rate above 0.5 Hz and assigned the excluded units the ranks of the next lowest ranked included unit. This was not done to exclude units from our analysis of the relation between spike waveform uniqueness and lifetime, but to group them more evenly. For all units with ranks smaller than a given percentile, we then estimated the conditional probability that a unit was lost in the subsequent session after having survived at least until that session (N). With *l*_*i*_ denoting the measured lifetimes of units, and Θ the Heaviside step function, that probability estimate was

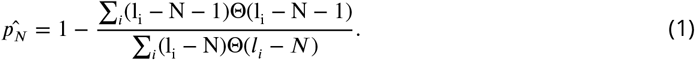

It assumes that after the N-th session, unit losses are described by a Poisson process with a fixed rate. The estimated additional lifetime (in sessions) *τ*_*N*_ was then given by

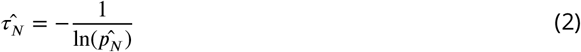

and shown in Figure 6 A. The same analysis (replacing ‘sessions’ by ‘days’) was performed to assess temporal lifetimes.

### Receptive fields

Firing rate responses were averaged across sessions and smoothed using a 41 ms Hanning kernel. Maximum responses were obtained for each stimulus condition and visualized. The receptive field was then determined as the region where the spatially interpolated response exceeded a threshold of twice the interquartile range above the median across conditions. Data were insufficient for estimating the size of the receptive field for individual sessions. To visualize the cross-session variation of receptive field locations, we assumed periodic boundary conditions and calculated the circular mean eccentricity and direction (colored circles in Figure 7 A, G). Temporal firing responses of individual sessions (Figure 7 D, J) were smoothed using an 18 ms Hanning kernel.

## Figure supplements

### Figure 4 — False discovery rate estimates

Spike shapes from different neurons can be similar, or even indistinguishable. To estimate how often we would falsely match a cluster from different units, we tried to match each cluster with clusters found on different channels within 3 sessions before and after its detection. The fraction of chance matches obtained from pairwise comparisons was then scaled by the number of clusters found on the same electrode to obtain an expected number of chance matches. This estimate assumes that the cluster would in fact be absent in the subsequent recording session (and would be lower otherwise). We further determined the dissimilarity threshold at which each pair of clusters would be matched to obtain a threshold dependence of the (pairwise) fraction of chance matches.

### Figure 4 — Statistics for long-term units

We determined variations in spike amplitude, rate and inter-spike intervals for long-term units by estimating relative standard deviations. For inter-spike intervals, specifically, we focused on short intervals, as these would more likely reflect intrinsic dynamics of a single neuron, rather than overall network behavior or stimulus dependent responses. In addition, these would also be more robust to potential contamination with noise.

We computed spike triggered spike count histograms in an interval from 0.2 - 50 ms after a spike. The first 0.2 ms were ignored as it would merely reflect noise in a few particularly noisy sessions, which were not the subject of this analysis. The histograms were converted into firing rates, smoothed using a 2 ms Hanning window, and normalized by the estimated firing rate of a given session, yielding an instantaneous, relative firing rate. Bursts of spikes would be reflected by an increased instantaneous firing rate shortly after a spike. For quantification, we measured the maximum of the instantaneous, relative firing rate, which was referred to as ‘burstiness’ in Figure 4 – Figure supplements 3 C and 4 C. As an estimate for a relative refractory period, we computed the temporal lag after a spike required to reach 3/4 of this maximum instantaneous firing rate.

As a summary statistic, we computed the fraction of the total variance across all clusters (from either group of long-term units), that the variation within units (and across sessions) could explain. This analysis was performed with logarithmized values in order to more equally weight clusters with lower averages.

### Figure 7 — Statistics for aggregate data

This analysis aimed at testing whether receptive field locations of identified units were consistent over time. Due to the retinotopic organization of area MT and the small size of the array, we expected similar receptive field locations across the array. Importantly, our sampling of space was relatively sparse and not perfectly homogeneous (few (i.e. 0-10) trials per condition). Additionally, there were few spikes per trial, as we analysed spiking in a short temporal window from 20 to 120 ms after stimulus onset.

To obtain a robust estimate of RF location with a high spatial resolution, we converted the sampled eccentricity and direction to unit vectors on a circle, to perform circular statistics (compute a resultant vector and compare to a uniform Poisson noise model). This approach may distort actual RF locations, but in the same manner for every dataset, and can therefore be used for comparing responses across sessions at a higher resolution. Specifically, we estimated receptive field locations by mapping the 5×7 grid of stimulus eccentricities and directions to circular variables equally spaced on unit circles. Summing up response vectors for different stimuli allowed forming a resultant vector with approximate multivariate Gaussian distribution for uniform responses (as null hypothesis), with a variance given by half the number of spikes in each of the 4 dimensions.

To account for different trial numbers for different conditions, we smoothed responses and trial numbers across directions and eccentricities using a [0.25 0.5 0.25] kernel (to ensure that there were no conditions without trials). We normalized each condition to reflect an average, per trial spike count and computed its variance under the assumption of probabilistic firing. Variances were then summed across conditions and divided by 2 (2 dimensions) to obtain an approximation of the variance of (each dimension of) the resultant vector under the null hypothesis. Comparing the resultant vector with the null hypothesis yields two numbers: (1) a sensitivity index, specific for a given receptive field location and independent of the number of trials. When treating the null hypothesis as a noise model and the resultant vector as the signal; both would have a variance of half the number of spikes, and hence the sensitivity index would be the length of the resultant vector divided by the square root of half the number of spikes. To obtain a sensitivity index independent of the number of trials, spike counts and resultant vectors were averaged across trials, allowing to compare individual sessions with the cross-session average.

(2) a p-value for accepting the null hypothesis of no spatial modulation. The half squared length of the resultant vector, divided by the total number of spikes is Chi-squared distributed with 4 degrees of freedom under the null hypothesis. Computing percentiles yielded p-values for each session.

It is a curiosity that units with larger sensitivity indices (Figure 7 A,B, red) tended to have receptive fields closer to the center of the region of detected receptive fields from the population than units with lower sensitivity indices (Figure 7 A,B, blue). We do not have an explanation for this observation, and neither did we have the statistical power to examine it in more detail.

## Acknowledgments

This work was supported by the US BRAIN Initiative (U01 NS094330) to ACH, the University of Texas at Austin (College of Natural Sciences Catalyst Award) to ACH, the National Institute on Drug Abuse (T32 DA018926) to AJL and HCC and the National Eye Institute (T32 EY021462) to AJL. We thank John P. Liska for comments on the manuscript.

## Competing interests

The authors declare that no competing interests exist.

---

**Figure 4–Figure supplement 1.**
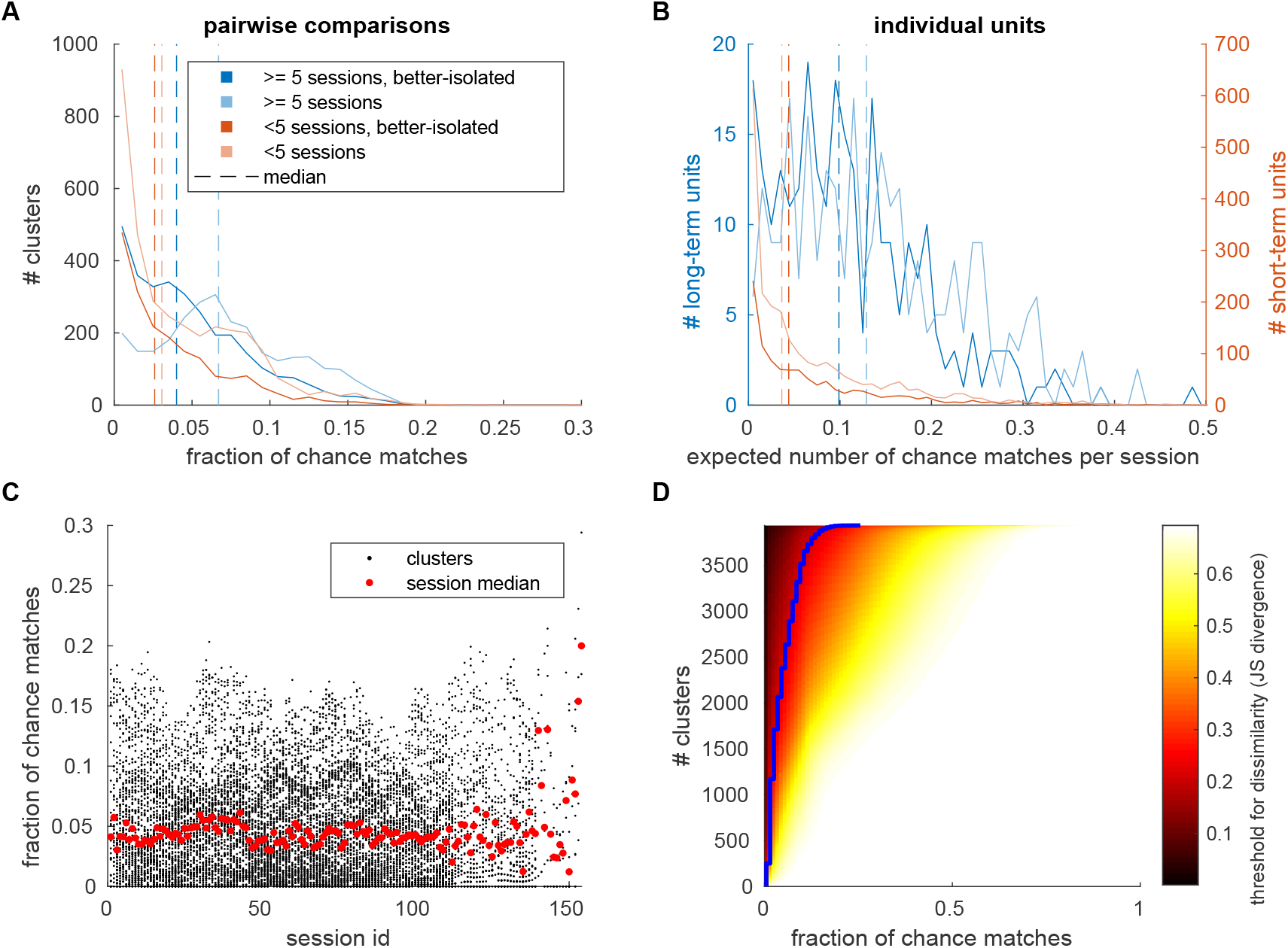
False discovery rate estimates for marmoset J. Spike shapes from different neurons can be similar, or even indistinguishable. To estimate how often we would falsely match a cluster from different units, we tried to match each cluster with clusters found on different channels within 3 sessions before and after its detection. (A) Histograms of the fraction chance matches in pairwise comparisons. Units were classified as in Figure 4 and the corresponding histograms were colored accordingly. Dashed lines mark median values. (B) Histograms of the average expected number of chance matches per session, when accounting for the number of detected clusters on the same electrode. (C) Pairwise false discovery rates across recording sessions. Red dots depict median values for each session. (D) False discovery rates in dependence of the dissimilarity threshold (blue line depicts threshold used in this work). Clusters were sorted according to the fraction of chance matches when using a fixed threshold.

**Figure 4–Figure supplement 2.**
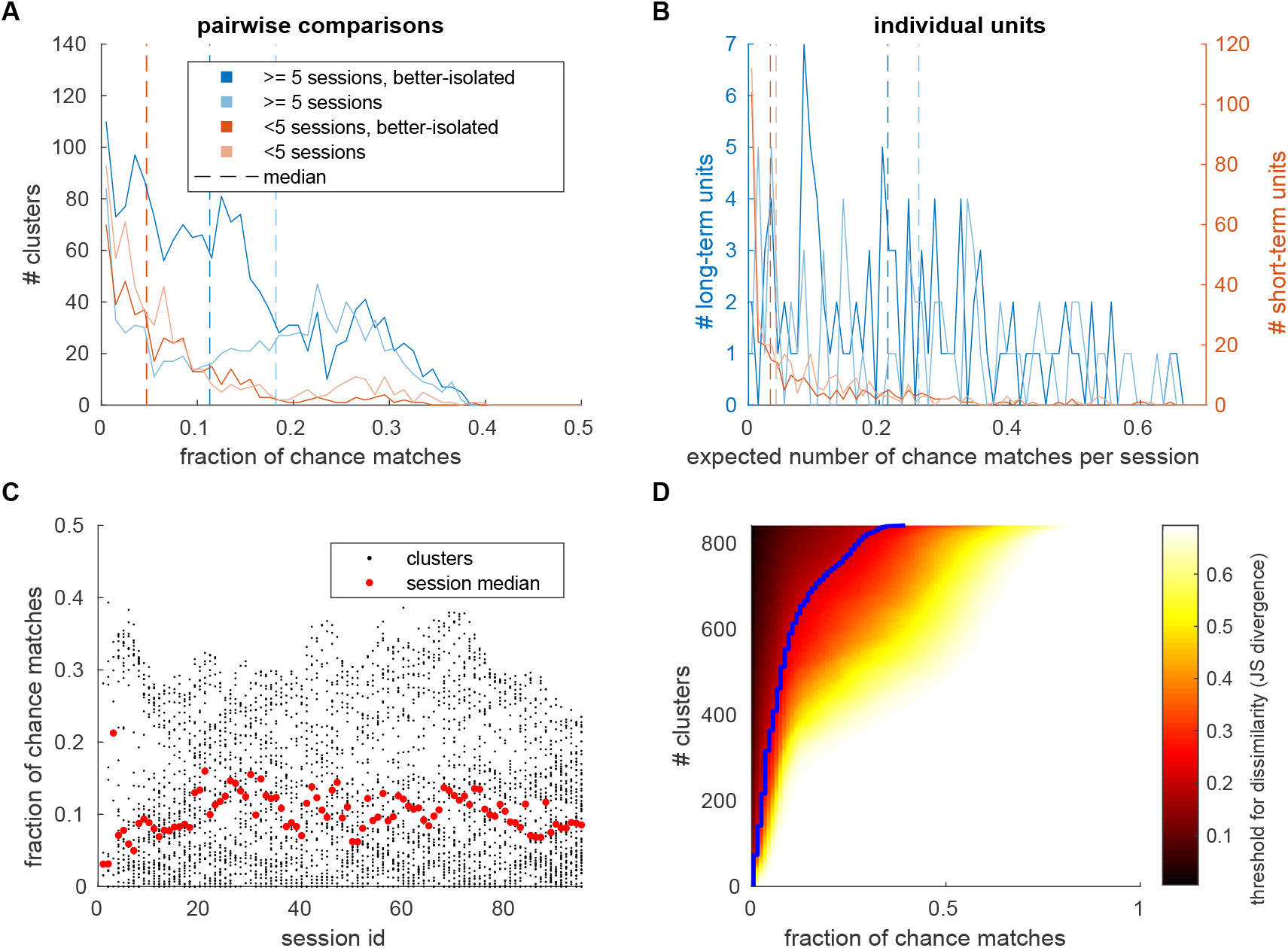
False discovery rate estimates for marmoset B. Spike shapes from different neurons can be similar, or even indistinguishable. To estimate how often we would falsely match a cluster from different units, we tried to match each cluster with clusters found on different channels within 3 sessions before and after its detection. (A) Histograms of the fraction chance matches in pairwise comparisons. Units were classified as in Figure 4 and the corresponding histograms were colored accordingly. Dashed lines mark median values. (B) Histograms of the average expected number of chance matches per session, when accounting for the number of detected clusters on the same electrode. (C) Pairwise false discovery rates across recording sessions. Red dots depict median values for each session. (D) False discovery rates in dependence of the dissimilarity threshold (blue line depicts threshold used in this work). Clusters were sorted according to the fraction of chance matches when using a fixed threshold.

**Figure 4–Figure supplement 3.**
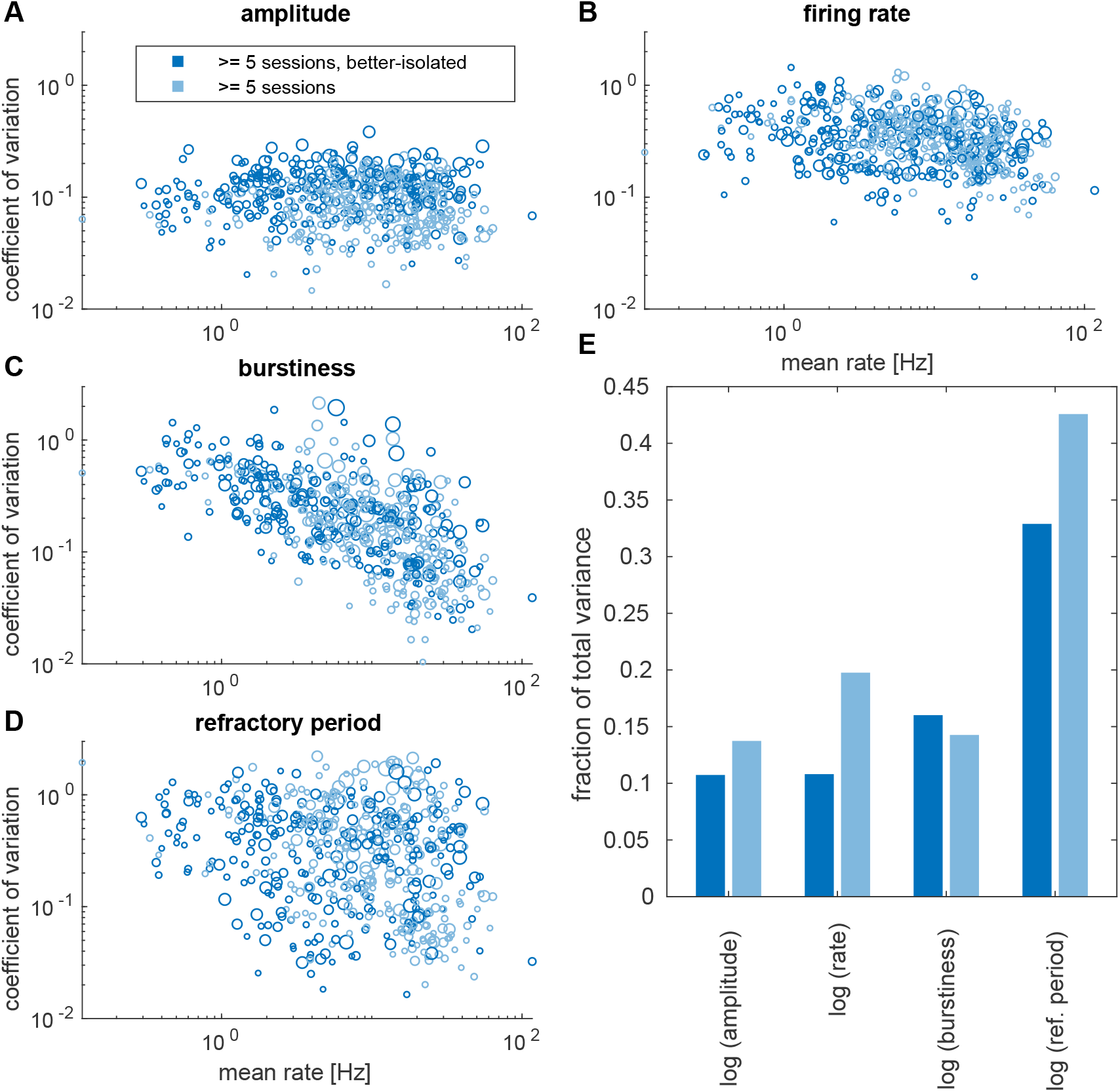
Long-term statistics for marmoset J. (A) Relative amplitude variations of long-term clusters. Larger symbols represent clusters observed in more experimental sessions, darker shades correspond to better-isolated units (as in Figure 4). (B) Relative firing rate variations. (C-D) Averages and variability of relative spike triggered averaged firing rates. To quantify the propensity of spiking in a short window after a spike, we computed spike triggered spike count histograms in an interval from 0.2 - 50 ms after a spike. These were converted into firing rates, smoothed using a 2 ms Hanning window, and normalized by the estimated firing rate of a given session. The maximum relative spike triggered firing rate was termed ‘burstiness’, and its variability for individual units is shown in (C). A high value would correspond to an increased chance of firing shortly after a spike, and a value around one would reflect no burst firing. As an estimate for a relative refractory period (variability shown in (D)), we computed the temporal lag after a spike required to reach 3/4 of this maximum firing rate. (E) Fraction of the total variance explained by within unit and across session variability. In order to more equally weight clusters with lower averages, this analysis was performed on a logarithmic scale.

**Figure 4–Figure supplement 4.**
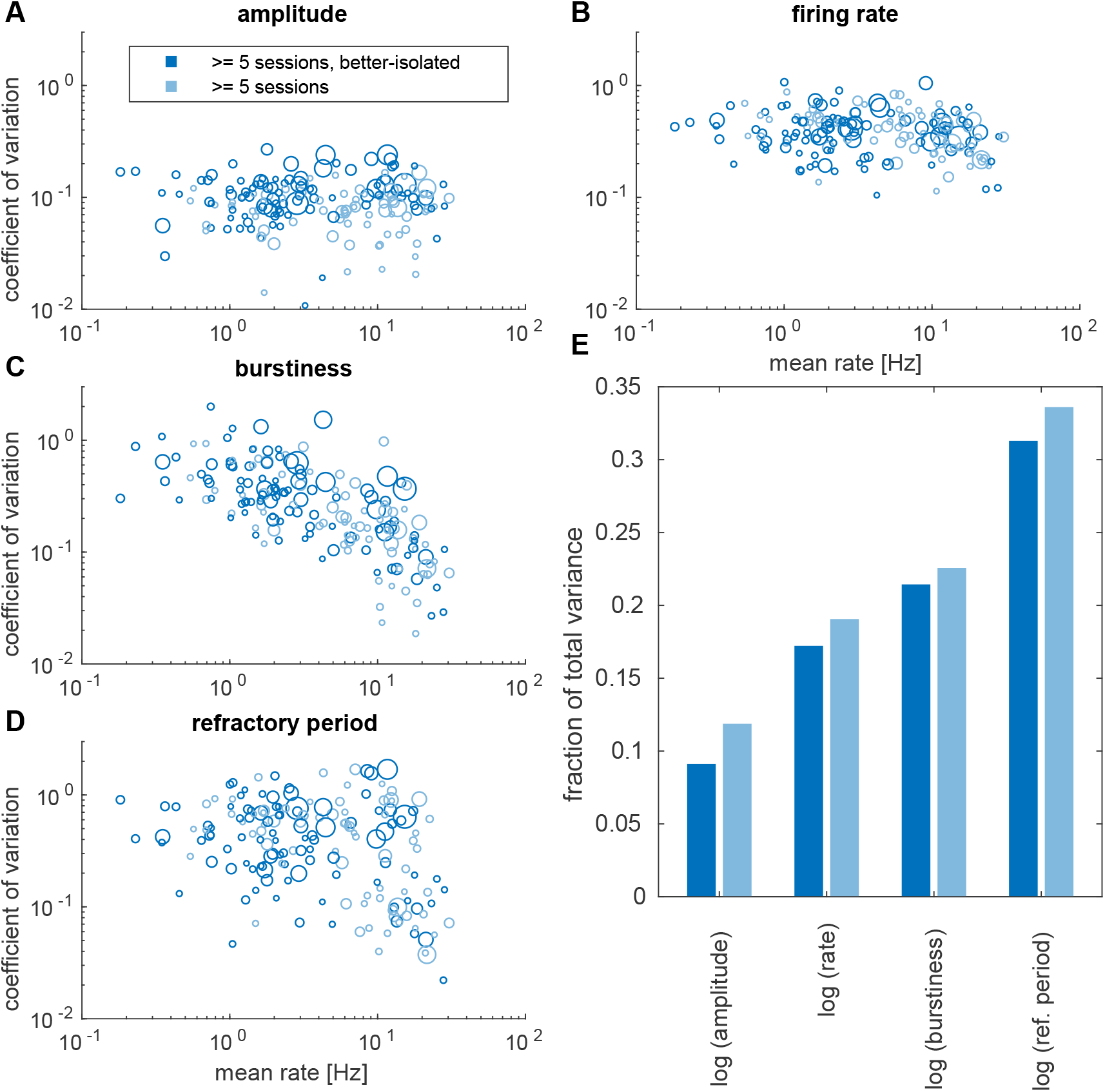
Long-term statistics for marmoset B. (A) Relative amplitude variations of long-term clusters. Larger symbols represent clusters observed in more experimental sessions, darker shades correspond to better-isolated units (as in Figure 4). (B) Relative firing rate variations. (C-D) Averages and variability of relative spike triggered averaged firing rates. To quantify the propensity of spiking in a short window after a spike, we computed spike triggered spike count histograms in an interval from 0.2 - 50 ms after a spike. These were converted into firing rates, smoothed using a 2 ms Hanning window, and normalized by the estimated firing rate of a given session. The maximum relative spike triggered firing rate was termed ‘burstiness’, and its variability for individual units is shown in (C). A high value would correspond to an increased chance of firing shortly after a spike, and a value around one would reflect no burst firing. As an estimate for a relative refractory period (variability shown in (D)), we computed the temporal lag after a spike required to reach 3/4 of this maximum firing rate. (E) Fraction of the total variance explained by within unit and across session variability. In order to more equally weight clusters with lower averages, this analysis was performed on a logarithmic scale.

**Figure 7–Figure supplement 1.**
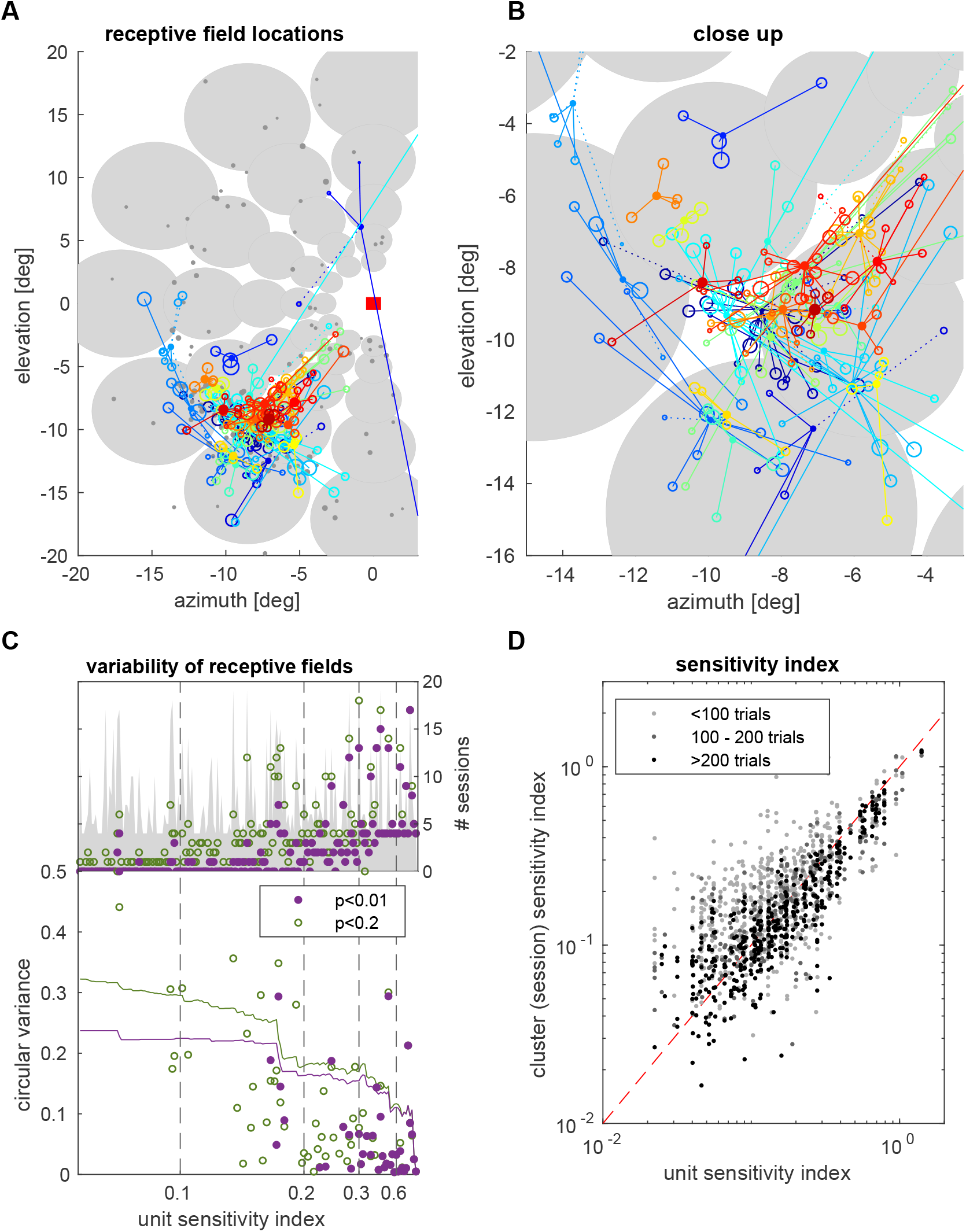
Statistics for aggregate data. Receptive field locations were estimated by mapping the 5×7 grid of stimulus eccentricities and directions to circular variables equally spaced on unit circles. Summing up response vectors for different stimuli allowed forming a resultant vector with approximate Gaussian distribution for uniform responses (as null hypothesis), and mapping the preferred stimulus location back to world coordinates. (A,B) Receptive field locations of units observed for at least 4 sessions with receptive field mapping (filled circles). Size/color relates to sensitivity indices (red: high, blue:low, gray:<0.3). Open circles denote estimated receptive field locations in individual sessions, linked to the corresponding unit with a solid line for sessions with a significant (p<0.01) spatial modulation of firing rates and and dotted line for a tendency (p<0.2) of a spatial modulation. (C) Variation of receptive field locations across at least 4 sessions from the same unit with good (p<0.01, purple) and weak (p<0.2, green) spatial modulation, normalizing individual session resultant vectors and computing the circular variance across sessions. The circular variance of a population of clusters from units with a given minimum sensitivity index is shown as a reference (colored lines). (D) Scatterplot comparing sensitivity indices of units computed across sessions and the corresponding single session estimates.

## Notes

### Competing Interest Statement

The authors have declared no competing interest.

